# BATMAN: fast and accurate integration of single-cell RNA-Seq datasets via minimum-weight matching

**DOI:** 10.1101/2020.01.22.915629

**Authors:** Igor Mandric, Brian L. Hill, Malika K. Freund, Michael Thompson, Eran Halperin

**Affiliations:** Department of Computer Science, University of California Los Angeles, Los Angeles, CA, USA; Department of Human Genetics, David Geffen School of Medicine, University of California Los Angeles, Los Angeles, CA, USA; Department of Anesthesiology and Perioperative Medicine, David Geffen School of Medicine, University of California Los Angeles, Los Angeles, CA, USA; Department of Computational Medicine, David Geffen School of Medicine, University of California Los Angeles, Los Angeles, CA, USA; Bioinformatics Interdepartmental Program, University of California Los Angeles, Los Angeles, CA, USA

## Abstract

Single-cell RNA-Sequencing (scRNA-Seq) is a set of technologies used to profile gene expression at the level of individual cells. Although the throughput of scRNA-Seq experiments is steadily growing in terms of the number of cells, large datasets are not yet commonly used due to prohibitively high costs. Integrating multiple datasets into one can improve power in scRNA-Seq experiments, and efficient integration is very important for downstream analyses such as identifying cell-type-specific eQTLs. State-of-the-art scRNA-Seq integration methods are based on the mutual nearest neighbors paradigm and fail to both correct for batch effects and maintain the local structure of the datasets. In this paper, we propose a novel scRNA-Seq dataset integration method called BATMAN (BATch integration via minimum-weight MAtchiNg). Across multiple simulations and real datasets, we show that our method significantly outperforms state-of-the-art tools with respect to existing metrics for batch effects by up to 80% while retaining cell-to-cell relationships. BATMAN is available at https://github.com/mandricigor/batman.

## 1 Introduction

Single-cell RNA sequencing (scRNA-Seq) has revolutionized transcriptomics as it enables the computational inference of cell types, the discovery of new cell states, and the reconstruction of cellular differentiation trajectories^1^. Although the scale of the datasets produced by scRNA-Seq is continuously growing^2^, the demand for sequencing an even larger number of cells greatly exceeds the current throughput of sequencing experiments^3^ and, therefore, cells must be processed in multiple sequencing runs, or batches. Due to the high level of technical noise, systematic differences between sequencing instruments, and other confounding factors, simple concatenation is a suboptimal approach to integrate multiple batches of a dataset. Further, batch effects have been shown to cause an increased number of false positives in downstream analyses^4^. To mitigate the level of false discoveries, a proper integration of multiple batches must eliminate the differences caused by batch effects.

Current methods for merging scRNA-Seq datasets can conventionally be divided into two categories: batch correction and integration methods^5^. Batch correction is the adjustment of gene expression in the high-dimensional gene space to account for confounding variation between technical scRNA-Seq replicates - in other words, batch correction operates on the gene expression levels themselves. Integration methods operate instead in a latent low-dimensional space, such as canonical correlation analysis (CCA) embeddings or embeddings learned from neural networks^6^, and are applied to the problem of merging multiple datasets across different technologies or biological conditions. Batch correction methods are more interpretable since they allow for a wider range of downstream analyses including differential gene expression and pseudo-time trajectory inference. On the other hand, integration methods enjoy a limited spectrum of applications, the most frequently used being visualization and cell-type classification. Throughout this paper, we will use the term “integration” for combining scRNA-Seq datasets into one as it is more general.

In this paper, we present a method for integration of single-cell datasets called BATMAN (BATch integration via minimum weight MAtchiNg), based on a parsimonious one-to-one matching of representative cells across datasets that maintains local structure. BATMAN operates in the high-dimensional gene space and uses the minimal total correction necessary to remove discrepancies between datasets. We show that BATMAN significantly outperforms state-of-the-art tools in terms of widely-used integration quality metrics on a wide range of simulated and real datasets (by 80%), but also maintains the local structure of each dataset.

## 2 Background

Given two scRNA-Seq datasets, a query dataset *D*_1_ and a reference dataset *D*_2_, the goal of integration is to align the query to the reference dataset by removing the confounding variation between them^7^. Usually after alignment only the query dataset is modified, i.e., the gene expression values of the cells in *D*_1_ are modified while the cells in *D*_2_ remain intact. The quality of the alignment is generally measured by a metric of how well-mixed the two cell populations are after modification. *D*_1_ and *D*_2_ are considered well-mixed if the local dataset-label distribution in the neighborhood of each cell (in the context of the integrated dataset *D*_3_) matches the global dataset label distribution, i.e, every ball containing cells of *D*_3_ contains cells from *D*_1_ and *D*_2_ in the same proportion as given by their cell counts^8^. The alignment is constrained by the local structure of each dataset; after integration, the neighborhood relationship among the cells in the query dataset must be preserved. If the datasets consist of several cell types, then the above definition applies to each cell type separately, i.e., cells of cell type *x* of dataset *D*_1_ should only be mixed with cells of cell type *x* of dataset *D*_2_.

scRNA-Seq dataset integration by maximizing the mixing quality of two datasets constrained by preserving their local structure is a challenging task as there are no straightforward or trivial objective functions to optimize over. Current state-of-the-art scRNA-Seq integration metrics such as kBET^8^ measure the quality of mixing through the concordance of the global and local dataset label distributions among the nearest k neighbors. Using such metrics as objectives in the integration problem prevents any standard optimization techniques from being applicable. Although it is possible to find a perfect mixing with respect to the aforementioned metrics by random assignment of cells in the dataset *D*_1_ to the cells of dataset *D*_2_, the biological signal of each dataset would be compromised as the local structure of each dataset would be destroyed.

One approach for solving the integration problem for scRNA-Seq datasets is to use tools designed for bulk RNA-Seq data. In this case, each cell in an scRNA-Seq dataset is viewed as a single bulk RNA-Seq sample. Nonetheless, integration methods that are borrowed from bulk RNA-Seq analysis depend on normalization techniques and usually assume that the data comes from a particular distribution. For example, ComBat^9^ and limma^10^ assume that the observed expression values come from a Gaussian distribution. These assumptions may not hold true for some real datasets. Additionally, ComBat and limma were not designed to handle datasets consisting of multiple cell types^11^. As it has been previously shown, in more complicated scenarios characteristic to single-cell datasets, these methods perform poorly^11^.

Many methods have recently been proposed for both integration and batch correction of scRNA-Seq data, primarily based on clustering or deep neural networks, for example, BERMUDA^11^, SAUCIE^12^, and Harmony^13^. BERMUDA uses a deep neural network architecture called an autoencoder to learn a low-dimensional representation of the data in an unsupervised manner. The autoencoder network is trained to minimize the loss between the original gene expression values and the reconstructed values after performing the dimensionality reduction, as well as the maximum mean discrepancy (MMD) between similar clusters from different batches. Like BERMUDA, SAUCIE also uses a sparse autoencoder to learn a low-dimensional representation of the data. The approach adds several novel regularization methods to the network activations to encourage the network to learn representations which are useful for several tasks in single-cell analyses such as clustering and integration. Harmony is an integration method that uses soft clustering after projecting the cells into a low-dimensional space by principal components analysis (PCA) to iteratively find cluster centroids, which are then used to calculate a cell-specific correction factor. The downside of these methods is that they operate in the latent space which limits their interpretability and use in downstream analyses such as differential gene expression and single-cell eQTL analyses.

Another group of integration methods that operate in the gene space includes Seurat v3.0^7^, MNNcorrect^14^, and Scanorama^5^. These methods can also be used for batch correction. MNNcorrect finds similar pairs of cells across batches where both cells are contained in each other’s set of nearest neighbors (mutual nearest neighbors, or MNN). The average difference in the gene expression between many pairs of mutual nearest neighbors estimates the batch effect, and this estimate can be used to correct the expression values. Seurat (version 3.0) builds on the MNN methodology, using MNN to determine “anchor points”. For dimensionality reduction, Seurat uses canonical correlation analysis (CCA) to find a subspace common to all datasets, which should be void of technical variation that is local to each dataset^7^,15. Scanorama also uses MNN for integration and batch correction, but the MNN search is performed in a low-dimensional space after randomized singular value decomposition (SVD) and uses a faster approximate nearest neighbor search to improve scalability. Additionally, Scanorama was designed to handle the alignment of multiple datasets without being sensitive to the ordering of alignment.

## 3 Parsimonious integration of scRNA-Seq datasets

Consider a query dataset *D*_1_ and a reference dataset *D*_2_. We formally define the batch effect vector for a cell:

### Definition.

*Suppose that a cell c is sequenced twice (i.e, the same cell sequenced in two different batches, D1 and D2) yielding two expression profiles* 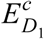 *and* 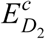. *The batch effect vector for cell c is defined as the vector* 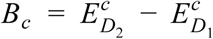.

One would be able to estimate batch effects between two datasets *D*_1_ and *D*_2_ if they shared a sample of cells. Indeed, according to the above definition, one would include RNA of a group of cells twice - once in *D*_1_ and once in *D*_2_ - compute batch effect vectors in these cells, and then extrapolate batch effect vectors onto the whole dataset. In reality, sequencing the same cell twice is infeasible (since the cell is destroyed in the sequencing process) and, therefore, it is impossible to directly compute the batch effect vectors of these cells. However, we postulate that if *D*_1_ and *D*_2_ originate from the same biological condition then for each cell *c*_1_ ∈ *D*_1_ there exists a cell *c*_2_ ∈ *D*_2_ such that the expression profile of 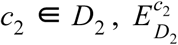 is closest to the expression profile of cell 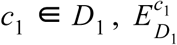 if *c*_1_ were to be sequenced twice (in *D*_1_ and *D*_2_). As we do not know which cell *c*_2_ ∈ *D*_2_ is closest to the unobserved expression profile 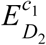, we search for such an alignment of *D*_1_ onto *D*_2_ which minimizes the total Euclidean distance between the cells in the dataset *D*_1_ and their corresponding (unobserved) cells in *D*_2_. In order to correct for batch effects, we need to compute the batch effect vector for each cell. Thus, we have to solve the following:

### Parsimonious Batch Effect Correction (PBEC) problem

*Given two scRNA-Seq datasets D*_1_ *and D*_2_, *find an alignment of the dataset D*_1_ *onto D*_2_ *for which the total length of the batch effect vectors across all the cells in D*_1_ *is minimized.*

The parsimony of batch effect vectors is strikingly intuitive. To illustrate the idea in a trivial case, suppose that we have two datasets and each dataset consists of two cells (Figure 1). There exist two different (one-to-one) alignments between the datasets. If we align the cells as depicted in Figure 1A, the total length of batch effect vectors is equal to 2 (the sum of the green edges). In this case, the local structure of the query dataset is preserved. However, if we align the cells as depicted in Figure 1B, then the total length of batch effect vectors is 4 and the local structure of the query dataset is compromised. This case only helps us illustrate the principle of the parsimonious alignment, and in order to emphasize its attractiveness over the existing methods we have to consider more challenging scenarios. MNN-based methods (in particular, MNNcorrect^14^) fail to properly correct for batch effects in cases when batch effects are not orthogonal to the biological signal or the magnitude of the batch effect vectors is large. Figures 2A and 2B show that parsimonious alignment provides a reasonable solution to the integration problem in these cases, while the mutual nearest neighbor based approaches fail.

**Figure 1:**
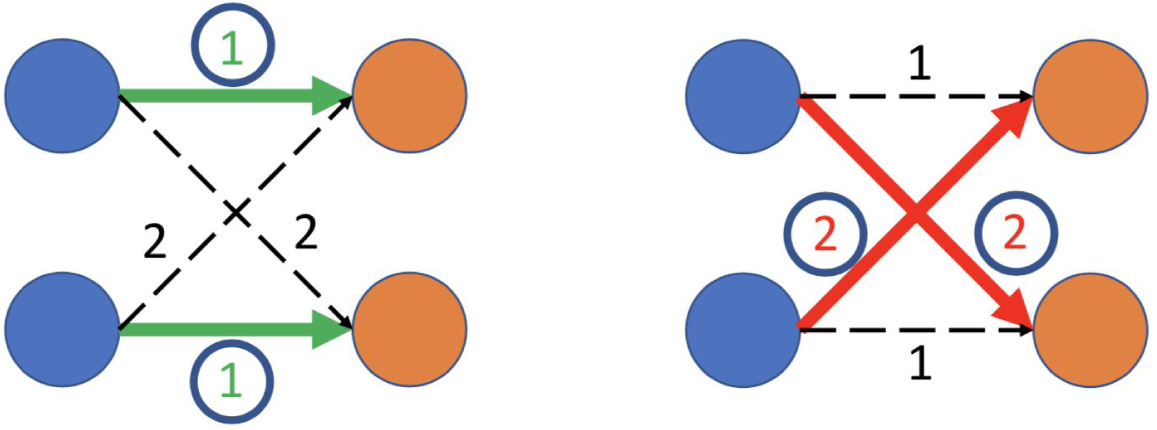
Possible anchor pairs. The two datasets (query - blue and reference - orange) consist of two cells. The weights on the edges are the Euclidean distances between the points. There are only two possible anchor pairs. A) The assignment is parsimonious since the total weight of the translation is equal to 2. The local structure of the query dataset is preserved. B) The assignment is not parsimonious (the total weight is 4) and it results in destroying the local structure of the query dataset.

**Figure 2:**
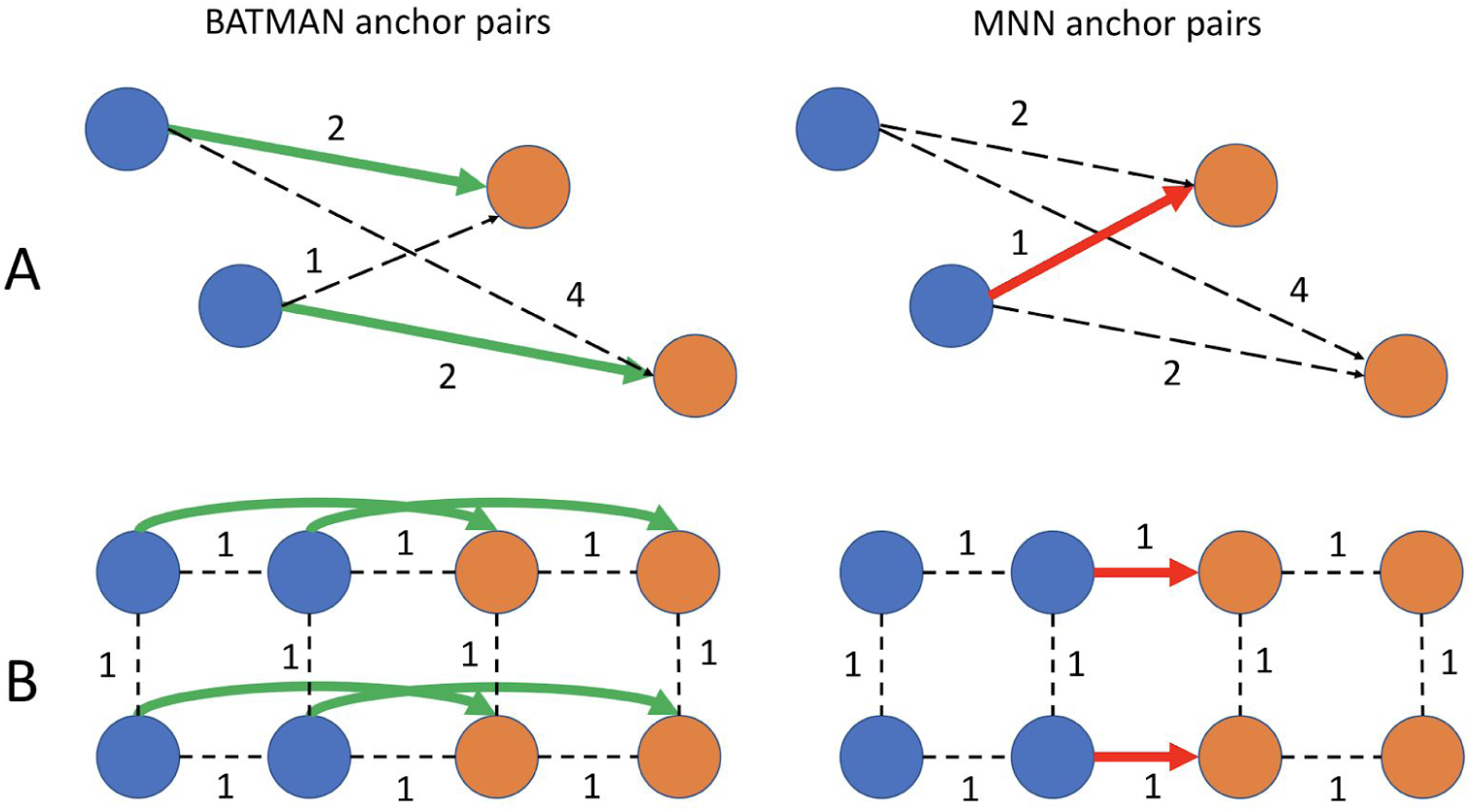
BATMAN anchor pairs vs mutual nearest neighbors anchor pairs. The edge weights are the Euclidean distances. A) The two datasets consist of two points each (blue and orange), the batch effects are **non-orthogonal** to the biological signal. BATMAN identifies the correct anchor pairs, while the MNN approaches fail. B) The two datasets consist of four points each and the batch effects are **large**. BATMAN identifies the correct anchor points, while the MNN approaches fail.

## 4 BATMAN: BATch integration via minimum weight MAtchiNg

In this section, we present our approach for solving the PBEC problem called BATMAN (BATch integration via minimum weight MAtchiNg). Suppose that we have two datasets, a query dataset *D*_1_ and a reference dataset *D*_2_. First, let us assume that |*D*_1_| = |*D*_2_ | and |*D*_1_| ≪ 1000. Then, solving the PBEC problem can be performed by computing the minimum weight matching in the weighted complete bipartite graph 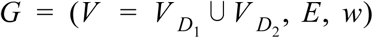 with *w*(*x, y*) = *d*(*x, y*), *x* ∈ *V* _1_, *y* ∈ *V* _2_, where *d*(*x, y*) is the Euclidean distance between gene expression profiles of the cell *x* in the query dataset and the cell *y* in the reference dataset. However, in practice, the equinumerosity of *D*_1_ and *D*_2_ is almost never met. Furthermore, the number of cells in a typical scRNA-Seq dataset is greater than 1000. Therefore, directly solving PBEC problem on large datasets with different number of cells is infeasible. To overcome this issue, we propose a novel algorithm, BATMAN. Instead of matching each cell in the query to a cell in the reference, BATMAN first identifies a set of representative cells in each dataset and then solves PBEC with respect to them. The solution of PBEC on the representative cells from *D*_1_ and *D*_2_ consists of a set of anchor pairs. The anchor pairs are then used to compute batch effect vectors in the representative cells. To determine batch effect vector in a cell belonging to *D*_1_ (not an anchor cell), we compute a weighted average of the batch effect vectors corresponding to the top *k* closest representative cells in *D*_1_. In more detail, BATMAN consists of the following steps:

1. ***Identification of representative cells.*** Representative cells of an scRNA-Seq dataset are the cells which are located in the high-density regions of the joint gene expression distribution. We propose finding representative cells by using clustering. As clustering in high-dimensional spaces is prone to multiple issues such as the “curse of dimensionality”^16^, we first compute PCA embeddings for each dataset separately. After, we perform clustering (for example, *K* -means: for small datasets up to 1000 cells, *K* ≈ 50; for larger datasets, *K* ≳ 300) on each of the two datasets and then identify the cluster centers *C*_1_ and *C*_2_ in the original gene space.
2. ***Building the anchor graph.*** We build the weighted complete bipartite graph *G* = (*C*_1_ ⋃ *C*_2_, *E, w*) - the *anchor graph*, where *C*_1_ and *C*_2_ are the cluster centers identified in the previous step of *D*_1_ and *D*_2_ respectively, and the weight of an edge (*x, y*) ∈ *E, x* ∈ *V* _1_, *y* ∈ *V* _2_ is equal to the Euclidean distance between the gene expression vectors *x* and *y*.
3. ***Computing the minimum weight matching.*** Next, we find the minimum weight bipartite matching in the anchor graph *G*. The endpoints of the edges belonging to the minimum weight matching represent the anchor pairs.
4. ***Computing the batch effect vectors in the representative cells.*** Based on the anchor pairs, we compute the batch effect vectors. Given an anchor pair (*x, y*), *x* ∈ *V* _1_, *y* ∈ *V* _2_, the corresponding batch effect vector for the cell *x* is the vector *T* _*x*_ = *y* − *x*.
5. ***Extrapolation of batch effect vectors and correction.*** For each cell *x* ∈ *D*_1_, we compute the batch effect vector *T* _*x*_ as a weighted average across the top *k* closest representative cells 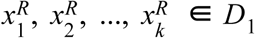:

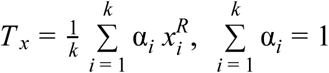

For cell *x*, we choose α_*i*_ to be proportional to 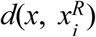. After the batch effect vectors *T* _*x*_ are determined in each cell *x* of *D*_1_, we correct for them, i.e. *x* → *x* + *T* _*x*_.

Dimensionality reduction of the datasets in Step 1 of our algorithm is crucial since it allows us to overcome the “curse of dimensionality” and reduces the runtime of clustering. The number of principal components should be large enough to ensure that the cells are separable by cell types (usually 20-30^7^.) Clustering ensures that the representative cells are distributed throughout the whole volume of the datasets (unlike for MNN-based methods; see Figure S1, Supplementary). Steps 2 and 3 represent the parsimonious alignment of the most representative cells between the two datasets which results in the set of anchor pairs. Finally, in Steps 4 and 5 we compute and correct for batch effects in each cell of the query dataset *D*_1_.

### Integrating datasets with multiple cell populations

In the case when the two scRNA-Seq datasets consist of multiple cell types (cell populations), we forbid anchor pairs with different cell type labels. That is, we propose filtering out edges of the anchor graph that are unlikely to be present in the matching solution. To accomplish this, we rely on transcriptional signatures of cell types, capturing the set of genes upregulated in each cell type. Assuming transcriptional signatures are characteristic to each cell type regardless of the technology, batch effects, or any other differences and systematic biases, we expect the correlation between two transcriptional profiles belonging to the same cell type to be high and the correlation of two transcriptional profiles belonging to different cell types to be lower. We use this expectation to filter out unlikely edges in the anchor graph. Namely, if the correlation between gene expression vectors 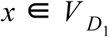 and 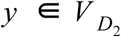 is below a threshold (for example, 0.7), then such an edge is removed from the graph. Filtering not only improves matching of the same cell types between the two datasets, but also significantly reduces the runtime of Step 4 of the BATMAN algorithm as the anchor graph becomes sparser.

### Non-concordant clustering between the datasets

Step 1 of BATMAN uses clustering to find the set of representative cells in each batch. A potential pitfall of such an approach is that in the case when the two datasets have different densities of their joint gene expressions as the naive minimum weight matching can fail due to the different sizes of the clusters being matched (Figure 3). In this case, the minimum weight matching can establish correspondences between clusters of significantly different sizes, and batch effects can not be fully corrected. To overcome this issue, we propose to replicate each representative cell according to the size of the cluster it belongs to. Namely, for larger clusters, we introduce additional representative cells (copies of the existing ones, Figure 3A). We make sure that the number of representative cells in the two datasets is the same. This helps to avoid biases caused by matching of representative cells whose clusters have significantly different sizes (Figure 3B).

**Figure 3:**
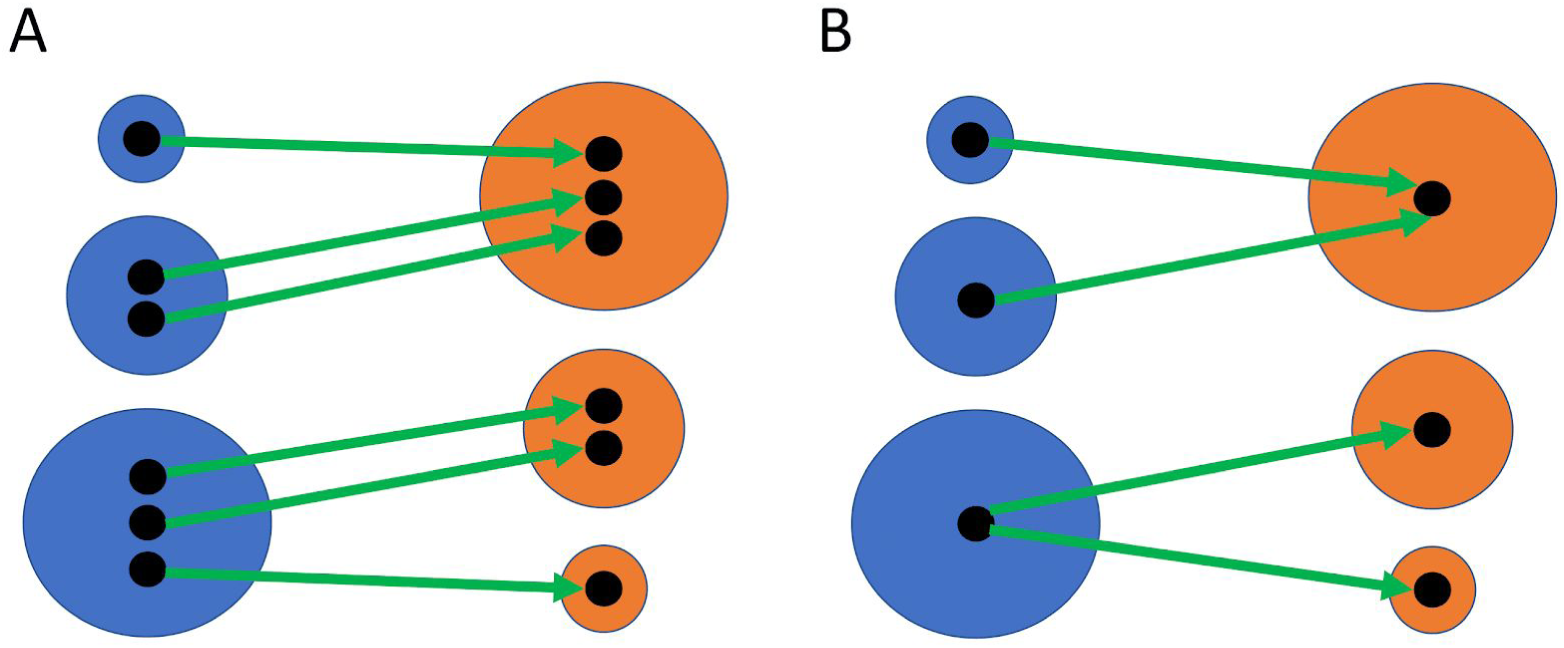
Non-concordant clustering. A) Introducing additional representative points; B) Final correspondence between clusters.

### Speeding up BATMAN

The steps of BATMAN are not computationally intensive except for Step 4, which involves the computation of the minimum weight matching in the anchor graph for a large enough number of anchors. Despite the fact that polynomial-time algorithms exist for its solving, it still represents a bottleneck for large graphs; for example, the well-known blossom algorithm has complexity *O*(*N* ^3^) ^17^. An optional speed-up which allows the application of BATMAN to large datasets uses approximation algorithms for the minimum weight matching. The standard greedy algorithm takes only *O*(*N log N*) time^18^ and, therefore, can be used to identify anchor pairs in very large graphs. However, it can yield a suboptimal solution which is at most twice as bad with respect to the total matching weight of the optimal solution.

## 5 Results

We compared BATMAN with three other state-of-the-art tools which operate in the gene space: Seurat v3.0, MNN, and Scanorama. We did not include tools such as limma and ComBat in the comparison, as they are are more appropriate for bulk RNA-Seq analysis and have been shown to fail in more complicated scenarios characteristic of single-cell datasets^11^. We also did not include SAUCIE, BERMUDA, and Harmony in the comparison since they operate on the latent spaces.

### Evaluation metrics

Traditionally, scRNA-Seq dataset integration quality has been assessed visually using UMAP and/or tSNE plots. However, there are multiple quantitative evaluation metrics available^7,11,14^. All of the metrics are based on scanning local neighborhoods of the cells in the combined dataset (i.e., after integration) and testing if the proportion of cells from the two datasets is the same as globally, for example, using the entropy mixing score. A novel test, kBET^8^, was designed independently of all the batch correction and integration methods and tests whether the local dataset label distribution is concordant with the global. As noted in (^11^) kBET has an issue: it fails to measure batch effects properly when the datasets have different cell-type compositions. Additionally, the more standard metrics such as entropy mixing score lack interpretability. To overcome these issues, we use LISI (Local Inverse Simpson Index), that is a novel recently proposed metric^13^. Its value ranges from 1 to 2 and it has a simple interpretation as the expected number of cells needed to be sampled before two are drawn from the same dataset. In our evaluations, we use two versions of LISI: integration LISI (iLISI; 1 is perfect separability, 2 is perfect mixing) and cell-type LISI (cLISI; 1 - all cell types are separable, 2 - cell types are mixed with each other).

To quantify how well a method preserves the local structure of datasets after integration, we measure the average percentage of retained nearest neighbors. Intuitively, if a cell and its neighbors are not perturbed by a batch correction method then they will have the same set of nearest neighbors after applying the correction. This quantity is computed as follows: given two datasets *D*_1_ and *D*_2_, for each cell we determine the top-*k* nearest neighbors (*k* -NN) in the original dataset. Next, for each cell we determine the top-*k* nearest neighbors after integration (again, in the context of its original dataset). We report the average percentage of retained nearest neighbors across all the cells in the union of *D*_1_ and *D*_2_. We will refer to this metric as *k* -RNN (retained nearest neighbors).

### Integration of simulated datasets

We simulated scRNA-Seq datasets based on a gamma-Poisson distribution using the *splatter*^*19*^ simulator (version 1.10.0) to compare BATMAN with Seurat V3.0, MNNcorrect (from scran R package, version 1.12.1), and Scanorama (version 1.5) across five different scenarios:

1. Large batch effects (LB)
2. Large batch effects with large dropout rate (LB-DR)
3. Large batch effects with unequal batch sizes (LB-UB)
4. Small batch effects (SB)
5. Large batch effects - two cell types (LB-CT)

Within Splatter, the magnitude of batch effects is controlled with two parameters: *batch.facLoc* and *batch.facScale*. Per Splatter’s documentation, for large batch effects scenarios in simulations we set both parameters to 0.5 and for small batch effects we set both to 0.001. The *dropout.shape* parameter controls the magnitude of dropout rate; setting this parameter to larger values produces sparser scRNA-Seq datasets. This parameter was set to 2 for the LB-DR scenario, and set to 1 in all other scenarios. In all five scenarios, we simulated 1000 genes. Batches in the LB-UB scenario consist of 200 and 1000 cells, while in the other scenarios each batch has 1000 cells. Finally, in the LB-CT scenario, each dataset consists of two cell types (80% and 20% cell type frequency in both batches).

For each scenario, we simulated 100 datasets and ran BATMAN alongside the other three tools. We computed the average values of iLISI and 50-RNN across the 100 runs and along with confidence intervals (CI) for each metric. The lower bound of the CIs were computed as the average of the 2nd and the 3rd value in the sorted list of 100 values, and the upper bound of the CIs were computed as the average of the 97th and the 98th values correspondingly. All the tools were run with their default parameters. In all the experiments, we used Seurat V3.0 pre-processing steps to obtain log-counts for each dataset.

In the LB scenario, a visual inspection of the datasets in the space of the top 2 principal components (See Figure 4) suggests that Seurat V3.0 slightly undercorrected for the batch effects. MNNcorrect and Scanorama, although correctly shifting the datasets towards the centers of each other, failed to properly account for the variance of the datasets as evidenced by Figure 4. However, BATMAN is the only tool that properly integrated two datasets; we show that the iLISI score of the integrated dataset is higher after integration only when applying BATMAN, and the other tools failed to properly correct for the differences between the datasets as evidenced by very low iLISI scores (see Table 1). For example, although the PCA plot of Seurat V3.0 shows that the batch effects were mostly removed (Figure 4), its low iLISI score suggests that in fact, it failed to properly integrate the two datasets. To investigate this discrepancy between the iLISI results and the PCA plots, for each integration result, we computed iLISI scores based on different numbers of top principal components (Figure 5A). When we consider top 2 principal components, the iLISI scores of Seurat V3.0 are high. However, as we increase the dimensionality of the dataset by projecting it to a larger number of principal components, the integration quality of Seurat V3.0 is constantly decreasing and in the limit it tends to 1. On the contrary, the original datasets look linearly separated in the top 2 principal components and the iLISI score is 1. However, when considering more top principal components, Figure 5A shows that in fact the two original datasets become better mixed with each other. BATMAN has the highest performance not only when considering the top 2 principal components, but also when considering more top principal components. It is the only method that manages to efficiently maintain a high iLISI score in a larger number of dimensions.

**Table 1:** Integration results in LB scenario (large batch effects): iLISI and 50-RNNscores. The best results are emphasized in bold. CI stands for confidence interval.

| Metric | Original | BATMAN | Seurat V3.0 | MNNcorrect | Scanorama |
| --- | --- | --- | --- | --- | --- |
| Mean iLISI | 1.82 | **1.84** | 1.00 | 1.00 | 1.01 |
| CI iLISI | (1.57, 1.92) | **(1.51, 1.96)** | (1.00, 1.02) | (1.00, 1.01) | (1.00, 1.01) |
| Mean 50-RNN | 1.00 | **0.92** | 0.58 | 0.51 | 0.14 |
| CI 50-RNN | (1.00, 1.00) | **(0.90, 0.95)** | (0.56, 0.61) | (0.51, 0.52) | (0.11, 0.16) |

**Figure 4:**
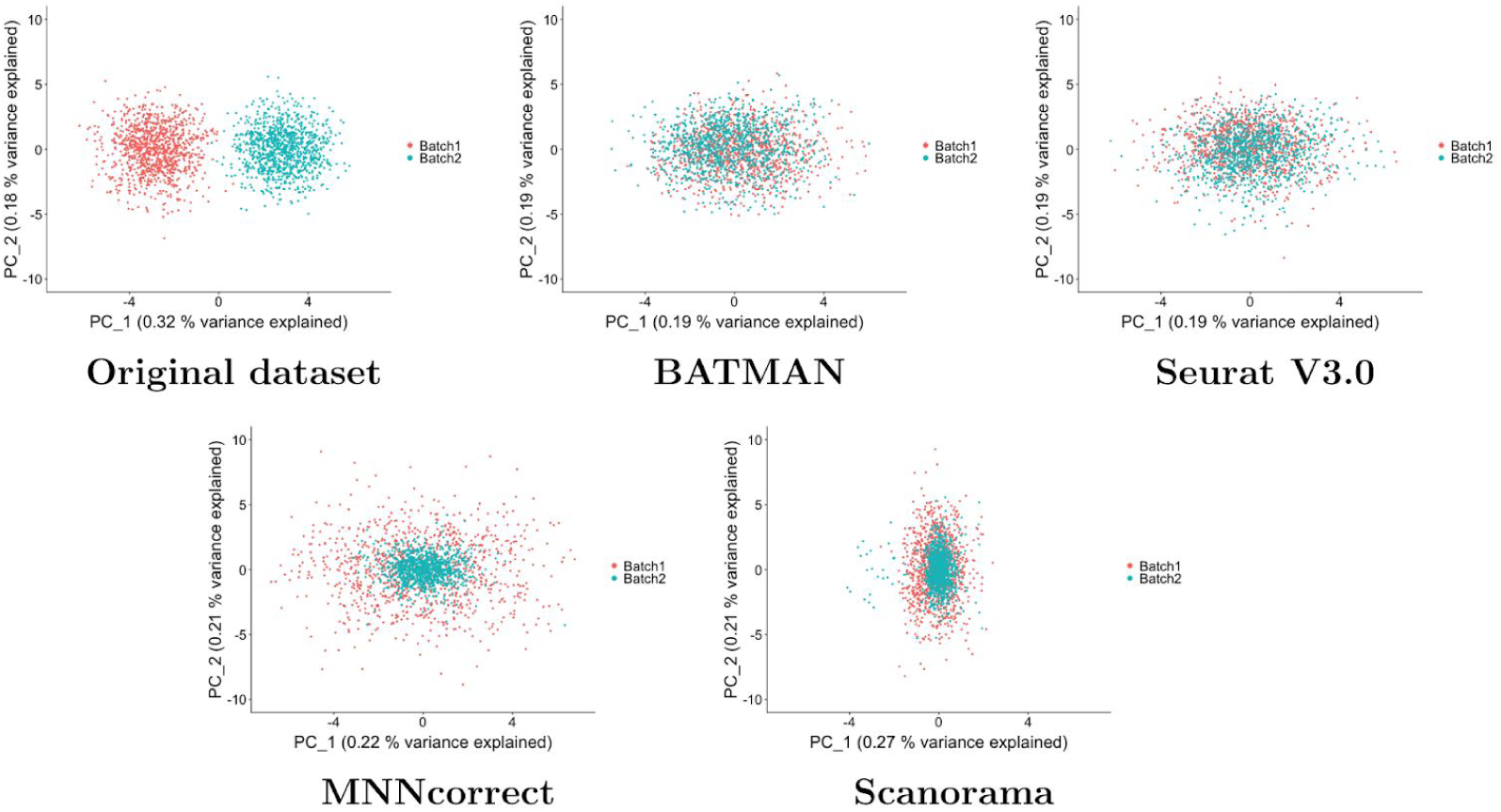
Simulated datasets with large batch effects (LB scenario) - PCA plots. Each dataset consists of 1000 cells and 1000 genes. The top 2 PCs are plotted.

**Figure 5:**
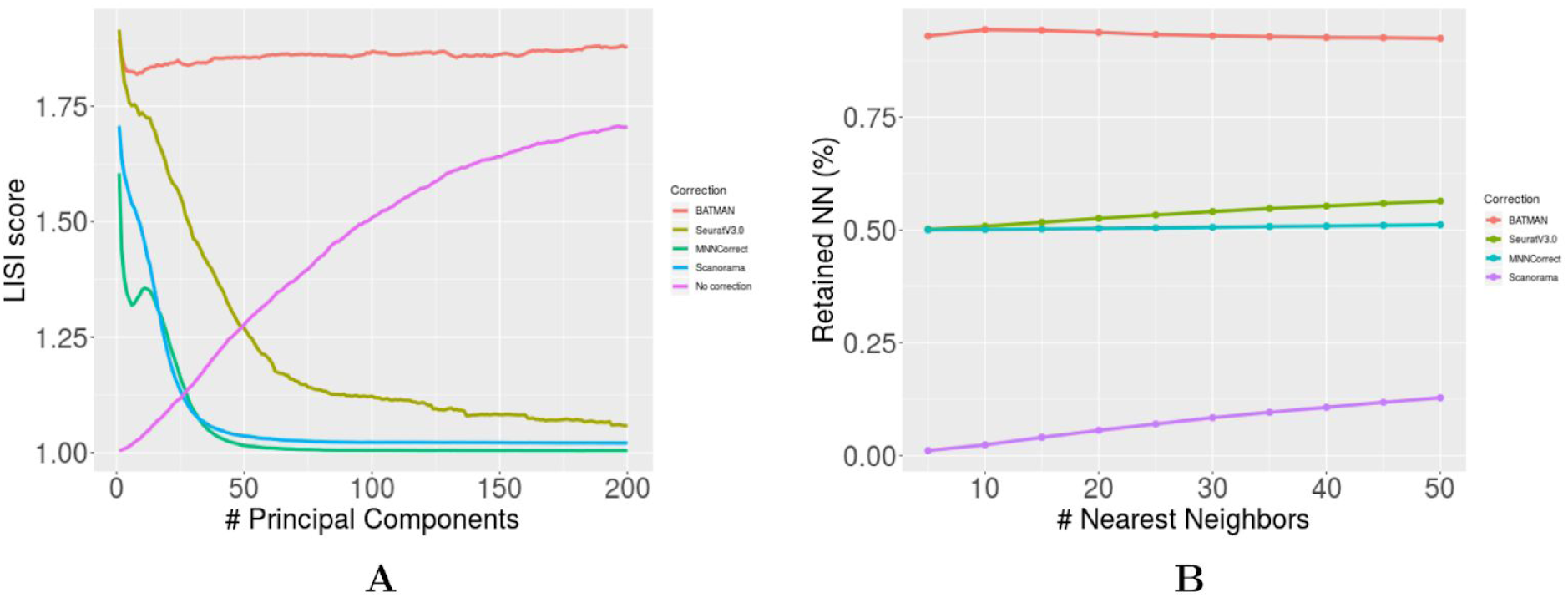
Evaluation of integration with large batch effects (LB scenario). A) iLISI score as a function of the number of top principal components. B) *k* -RNN score for different values of *k*.

Notably, BATMAN is the only tool that not only properly corrected for the batch effects but also preserved the biological signal characteristic to each dataset before integration (see Table 1). The 50-RNN of BATMAN equal to 0.92 means that on average, BATMAN preserves 92% of each cell’s 50 nearest neighbors after integration. Seurat V3.0 and MNN each preserve 58% and 51% correspondingly, while Scanorama integrates two datasets without keeping their local structure resulting in a poor 50-RNN score of 0.14. BATMAN, Seurat V3.0, and Scanorama display a consistent *k* -RNN score across multiple values of *k*, while Scanorama has very poor results for smaller values of *k* (reflecting that it destroys local structure in each small neighborhood of the two datasets; Figure 5B).

Higher dropout rates (the LB-DR scenario) do not seem to drastically change the results of integration across the four methods (Figure S2, Supplementary) as compared to the LB scenario. Again, BATMAN significantly outperformed the other tools on iLISI and 50-RNN scores (see Table S1, Supplementary).

When the two datasets consist of different number of cells (the LB-UB scenario), BATMAN is the only method demonstrating any improvement in the iLISI score after integration (see Figure S3, Supplementary and Table S2, Supplementary). Notably, both BATMAN and Seurat V3.0 preserved the local structure of the datasets with a 50-RNN score above 0.7 (see Figure S3, Supplementary and Table S2, Supplementary).

We also measured the ability of the four methods to properly correct for very small batch effects (the SB scenario). In this case, the two datasets were well-mixed even before integration with an initial iLISI score of 1.88. Given that any integration method will attempt to correct for batch effects even when the datasets are perfectly mixed, in this scenario a decreased iLISI score after integration indicates an overcorrection. As expected, the iLISI score for each of the four methods is smaller than the score before integration (see Table S3, Supplementary). However, the smallest overcorrection is achieved by BATMAN (iLISI = 1.87) while the other methods’ overcorrection introduced large batch effects.

Finally, when two cell types are simulated (the LB-CT scenario), MNNcorrect, BATMAN, and Scanorama achieve high iLISI scores (of at least 1.6) while Seurat V3.0 fails to properly integrate the two datasets. Interestingly, MNNcorrect slightly outperformed BATMAN both in terms of iLISI and 50-RNN scores (see Figure S5 and Table S4, Supplementary).

The analysis of iLISI and k-RNN scores in PCA embeddings suggest that in most of the simulation scenarios, MNN-based methods only work well in latent spaces of up to 100 principal components, while BATMAN successfully maintains a high quality of integration in higher dimensions. BATMAN also retains the local structure by preserving cell-to-cell relationships for each integrated dataset while the MNN-based methods perform worse (see Figures S6-S9, Supplementary).

### Integration of real datasets across different technologies

To compare BATMAN with the other tools in the case of real datasets across different single-cell technologies, we downloaded two pancreatic datasets: a CEL-Seq2 dataset consisting of 2285 cells^20^ and a Smart-Seq2 dataset consisting of 2394 cells^21^. Both datasets include 13 pancreatic cell types with different cell-type composition (see Table S5 for cell-type composition and Figure S10, Supplementary for UMAP plots).

To examine how different methods correct for batch effects and match cell type populations, we performed two experiments: (1) integration of full datasets (“all cell types”), and (2) integration with one cell type missing from one of the datasets (“1-held-out”). The second experiment evaluates whether the integration methods can align cells of the same cell types across datasets. For maximum stringency, we excluded the most abundant cell type (alpha cells, 37% of cells in CEL-Seq2 dataset and 42% of cells in Smart-Seq2 dataset - see Table S5, Supplementary) from CEL-Seq2 dataset.

Firstly, when all cell types are present across the two datasets, all four methods perform well on matching cell-type populations; the cell types are initially well-separable (cLISI score 1.05, Table 3) and they remain so after the integration of all four methods. However, there are large batch effects between the two datasets (iLISI score 1.07). Despite the fact that the integration results look successful upon visual inspection for all methods (see UMAP plots of Figure 6), BATMAN outperformed the MNN-based methods (iLISI of 1.55 for BATMAN versus next highest iLISI of 1.35 for Seurat V3.0).

**Table 3:** Integration results for real pancreas dataset. The best results (out of four methods) are marked in bold. For iLISI, higher correspond to higher quality of integration; for cLISI, lower scores correspond to better separation of cell types.

| Experiment | Score | No correction | BATMAN | Seurat V3.0 | MNNcorrect | Scanorama |
| --- | --- | --- | --- | --- | --- | --- |
| All cell types | iLISI | 1.07 | **1.55** | 1.35 | 1.18 | 1.22 |
|  | cLISI | 1.05 | 1.08 | **1.03** | 1.08 | 1.04 |
| 1-held-out | iLISI | 1.09 | **1.39** | 1.23 | 1.20 | 1.21 |
|  | cLISI | 1.04 | 1.07 | **1.03** | 1.09 | 1.05 |

**Figure 6:**
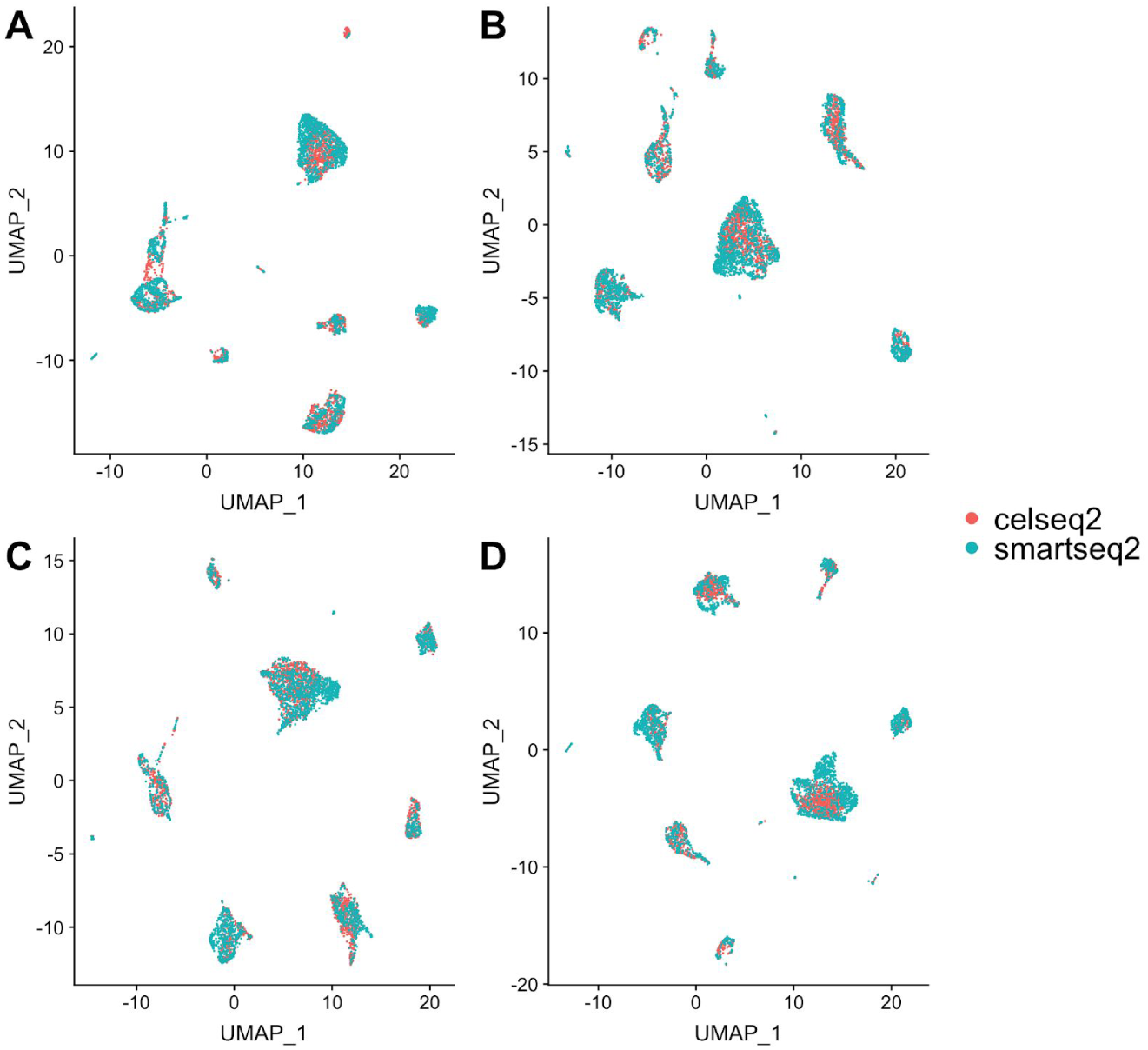
Integration of the two pancreatic datasets across different technologies - UMAP plots (“all cell types” scenario). A) BATMAN; B) Seurat V3.0; C) MNNcorrect; D) Scanorama.

Notably, BATMAN is outperformed by Seurat V3.0 and MNNcorrect in lower dimensions as iLISI scores for BATMAN are slightly worse than those for Seurat V3.0 and MNNcorrect in top-50 to top-170 principal components (Figure 7A). However, as the number of dimensions grows, BATMAN’s iLISI score increases and the iLISI scores of MNN-based methods decrease. BATMAN showed significantly higher *k* -RNN scores compared to the other methods (Figure 7B) which demonstrates that BATMAN better preserves the local structure of each dataset.

**Figure 7:**
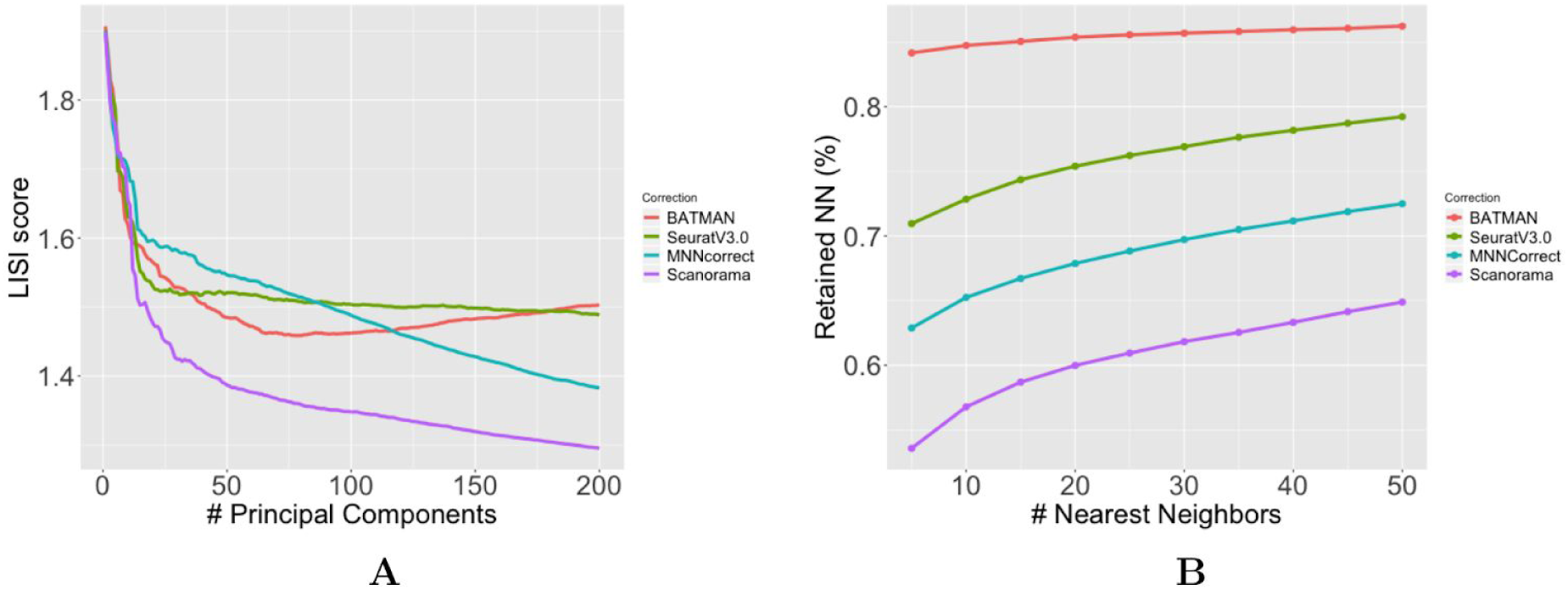
Integration of the two pancreatic datasets across different technologies: integration quality and local structure preserving. A) iLISI score as a function of top principal components; B) *k* -RNN metric across different values of *k*.

Secondly, in the 1-held-out experiment, all methods correctly matched the cell types leaving alpha cells from Smart-Seq2 dataset unmatched (Table 3 and Figures S12-S13, Supplementary). The cLISI scores of all the methods are approximately the same, while the iLISI scores vary. BATMAN achieves the best iLISI score (1.39), while the worst iLISI scores are produced by MNNcorrect and Scanorama integration results. Again, BATMAN introduced the least deformation to the local structure of the datasets compared to the MNN-based methods (Figure S14, Supplementary).

In terms of runtime, BATMAN and Scanorama perform best. In the “all cell types” experiment, BATMAN and Scanorama integrated the two datasets in 9 and 14 seconds, while Seurat V3.0 and MNNcorrect - in 22.3 and 228.6 seconds correspondingly. In “1-held-out” experiment, the runtimes were slightly smaller: BATMAN - 4 seconds, Scanorama - 11.8 seconds, Seurat V3.0 - 16.3 seconds, and MNNcorrect - 190.8 seconds. All the experiments were performed on a MacBook Pro laptop (16 GB RAM, 8 core 2.3 GHz Intel Core i9 processor).

### Integration of real 10X Genomics datasets

Current trends in single-cell RNA sequencing suggest that droplet-based technologies such as 10X Genomics will become the state-of-the-art in single-cell data acquisition due to their ability to simultaneously profile large numbers of cells in one experiment. Also, their cost-effectiveness makes them more attractive for large-scale experiments. Therefore, it is crucial for integration methods to be able to handle massive sparse datasets like those of 10X Genomics, which have a higher dropout rate.

We sought to assess the ability of BATMAN to integrate two 10X Genomics datasets with different numbers of cells and different cell type composition. For this purpose, we downloaded the PBMC 8k dataset (8381 peripheral blood mononuclear cells) and the “Pan T cells” dataset (3555 T cells) from 10X Genomics’ official website. The cells in the PBMC 8k dataset belong to multiple cell types such as CD4 T cells, B cells, dendritic cells, and more; thus, we expect the Pan T cells dataset to be biologically similar to a subset of the PBMC 8k dataset. Indeed, the PCA plots reveal that PBMC 8k dataset consists of 3 clusters (Figure 8A; “pbmc8k_0”, “pbmc8k_1”, and “pbmc8k_2”), while the Pan T cells dataset consists of a single cluster (Figure 8B; “t_3k”). On a PCA plot of the combined dataset without integration in Figure 8C, t_3k is closer to pbmc8k_0. The expression plots of two immune marker genes IL7R and NKG7 (Figure 8D) reveal a shared biological signal between the two clusters. However, large batch effects are present (Figure 8C).

**Figure 8:**
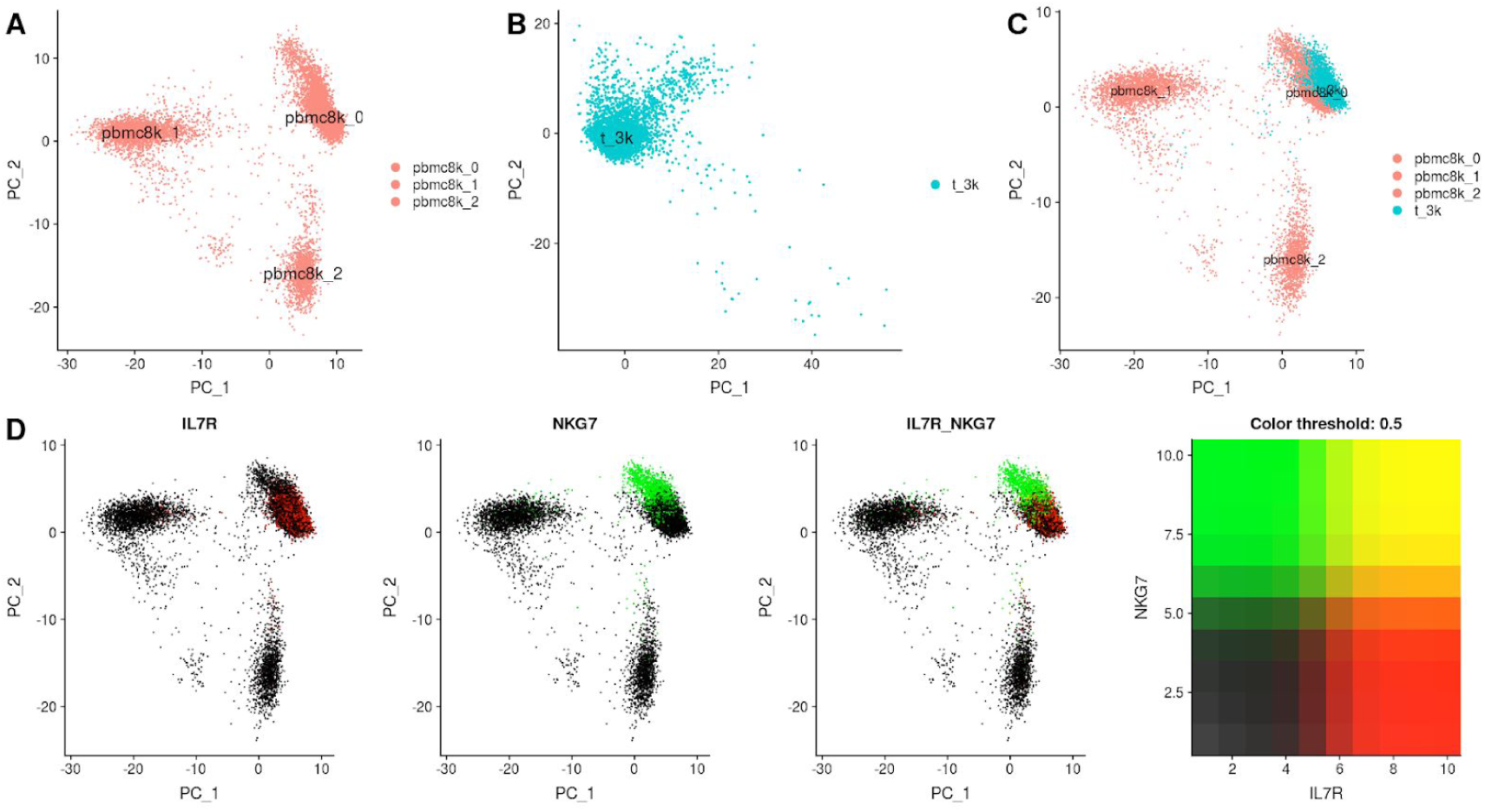
PBMC 8k and Pan T cells datasets (10X Genomics). A) PCA plot of PBMC 8k dataset reveals 3 clusters: pbmc8k_0, pbmc8k_1, and pbmc8k_2;. B) PCA plot of Pan T cells dataset reveals 1 cluster - t_3k. C) PCA plot of both datasets with no integration shows significant batch effects between cluster pbmc8k_0 of PBMC 8k dataset and t_3k cluster of Pan T cells dataset; D) Feature plot showing coexpression of IL7R and NKG7 marker genes. Pan T cells dataset corresponds to cluster pbmc8k_0 of PBMC 8k dataset. A proper integration of these two datasets should mix these two clusters.

We integrated the two datasets using BATMAN and the three MNN-based methods. BATMAN not only had the highest iLISI score of the four methods, but it also better preserved cell-to-cell relationships (Figure 9A). MNNcorrect performed slightly worse than BATMAN in terms of both integration and local structure preserving (Figure 9C). Although Scanorama had a good iLISI score, it destroyed much of the cell-to-cell relationships (Figure 9D). Finally, Seurat V3.0 failed to properly match the biological state of the cells between the two datasets (Figure 9B) which resulted in lower quality of integration.

**Figure 9:**
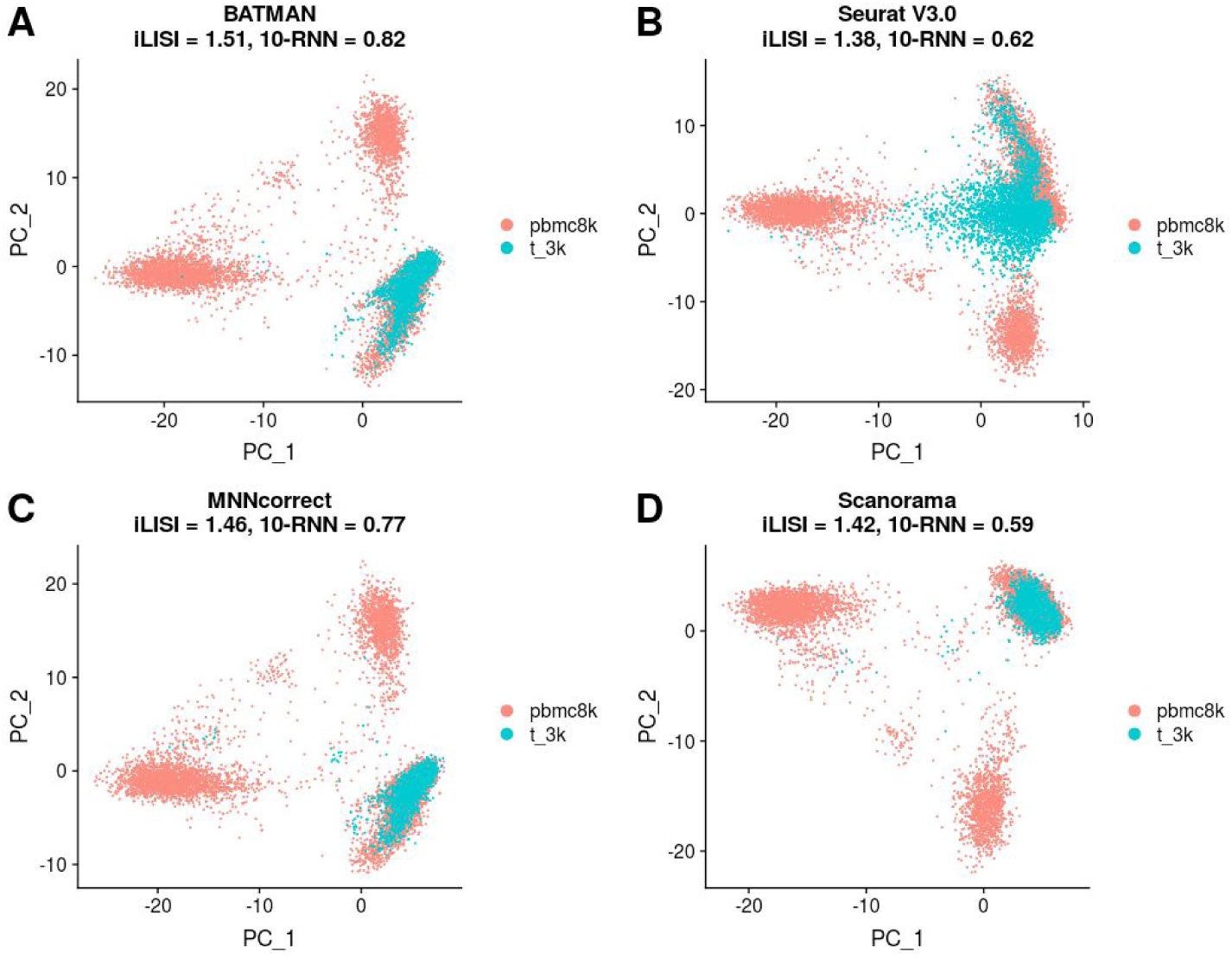
Integration of PBMC 8k and Pan T cells datasets (10X Genomics). A) BATMAN; B) Seurat V3.0; C) MNNcorrect; D) Scanorama.

In terms of runtime, BATMAN and Scanorama were the fastest tools finishing the integration task of the two 10X Genomics datasets in 20.1 and 31.0 seconds correspondingly. Seurat V3.0 used 211.2 seconds, while MNNcorrect was the slowest tool - 1099.5 seconds.

## 6 Discussion

We present BATMAN, a novel method for scRNA-Seq dataset integration. It shows significantly better performance than state-of-the-art methods for mixing two datasets while efficiently matching cell type populations and preserving the cell-to-cell relationships of each dataset. The underlying innovation of BATMAN is novel and has never been applied before in scRNA-Seq integration. Our parsimonious formulation based on minimum-weight matching not only finds a minimal correction needed to integrate two datasets, but also preserves the intrinsic structure of the two datasets.

We show that BATMAN achieves better performance than the state-of-the-art methods in terms of widely used metrics such as LISI on a wide range of datasets.

The current implementation of BATMAN works only with two datasets. However, in principle, it can be used to integrate multiple datasets (similar to multiple integration in MNNcorrect).

## Supplementary materials

**Figure S1:**
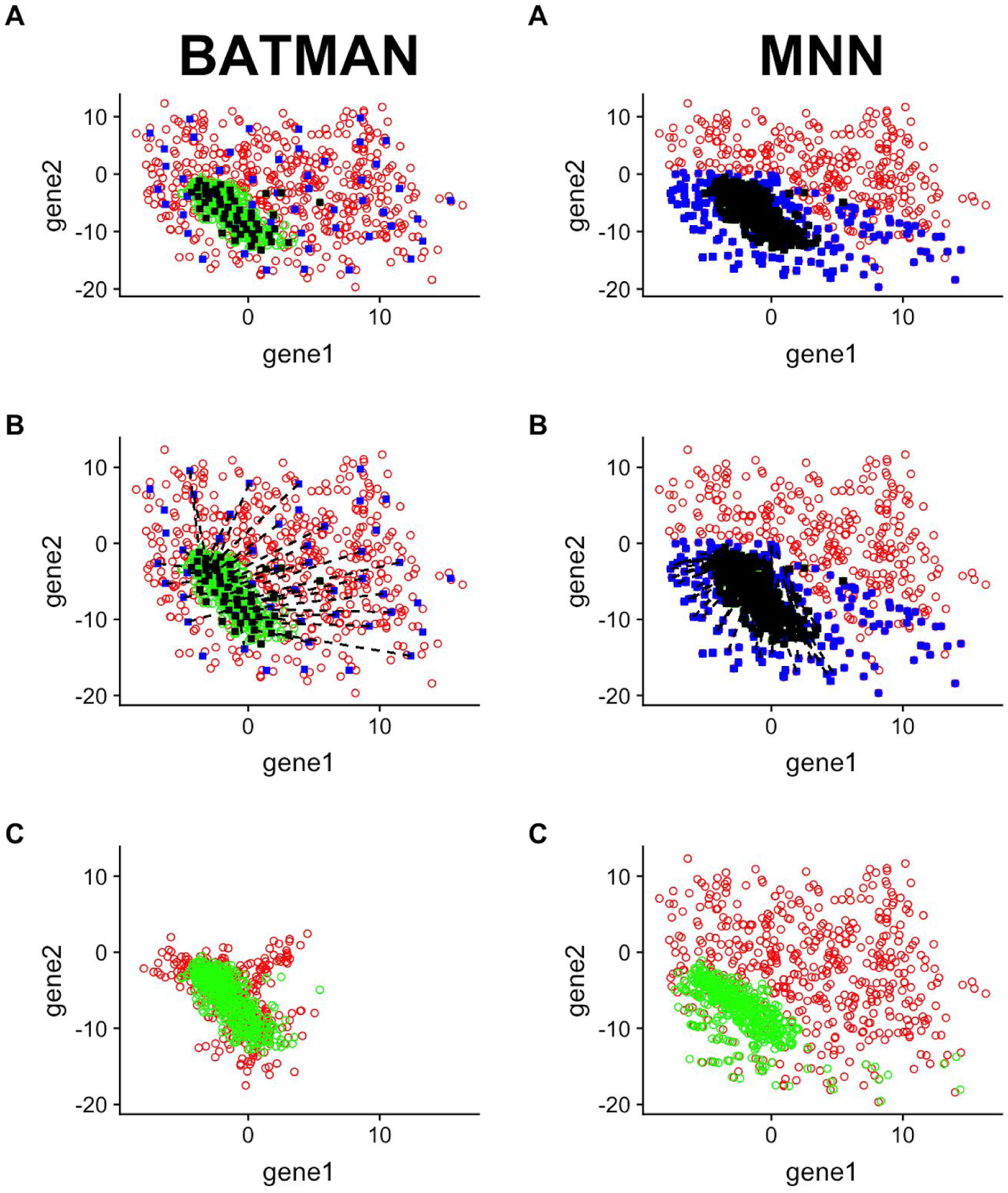
Choosing anchor cells. (A, B) BATMAN considers anchor cells across all the volume of both datasets, while MNNcorrect (and other MNN-based methods) consider only anchors at the frontier between the datasets. (C) BATMAN successfully integrates two datasets, while MNN-based methods fail.

**Figure S2:**
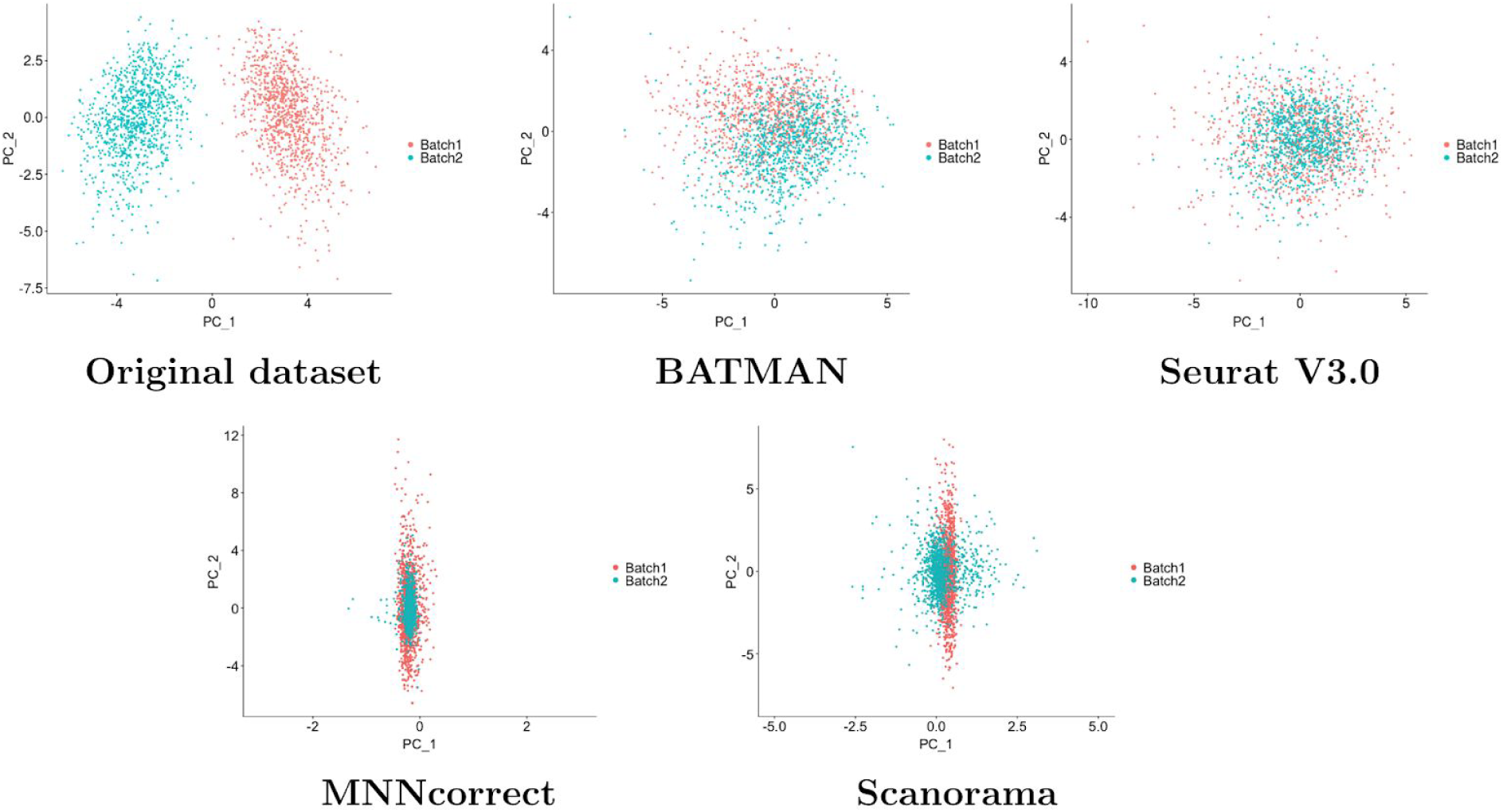
Simulated datasets with large dropout (LB-DR scenario) - PCA plots. Each dataset consists of 1000 cells and 1000 genes. The top 2 PCs are plotted.

**Table S1:**
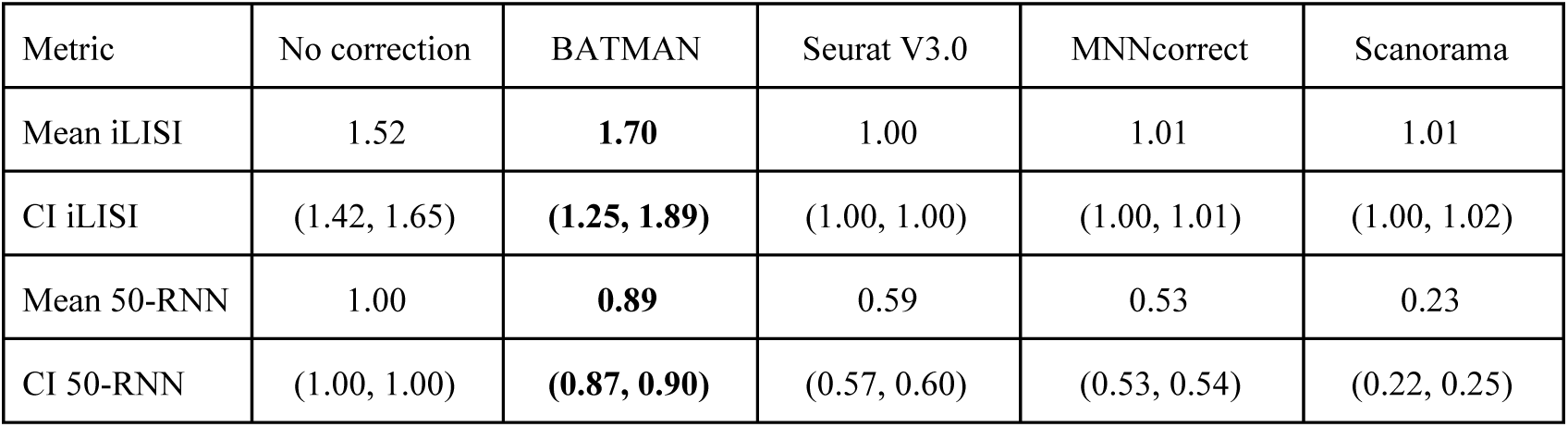
Integration results in LB-DR scenario (large dropout): iLISI and 50-RNNscores. The best results are emphasized in bold. CI means confidence interval.

| Metric | No correction | BATMAN | Seurat V3.0 | MNNcorrect | Scanorama |
| --- | --- | --- | --- | --- | --- |
| Mean iLISI | 1.52 | **1.70** | 1.00 | 1.01 | 1.01 |
| CI iLISI | (1.42, 1.65) | **(1.25, 1.89)** | (1.00, 1.00) | (1.00, 1.01) | (1.00, 1.02) |
| Mean 50-RNN | 1.00 | **0.89** | 0.59 | 0.53 | 0.23 |
| CI 50-RNN | (1.00, 1.00) | **(0.87, 0.90)** | (0.57, 0.60) | (0.53, 0.54) | (0.22, 0.25) |

**Figure S3:**
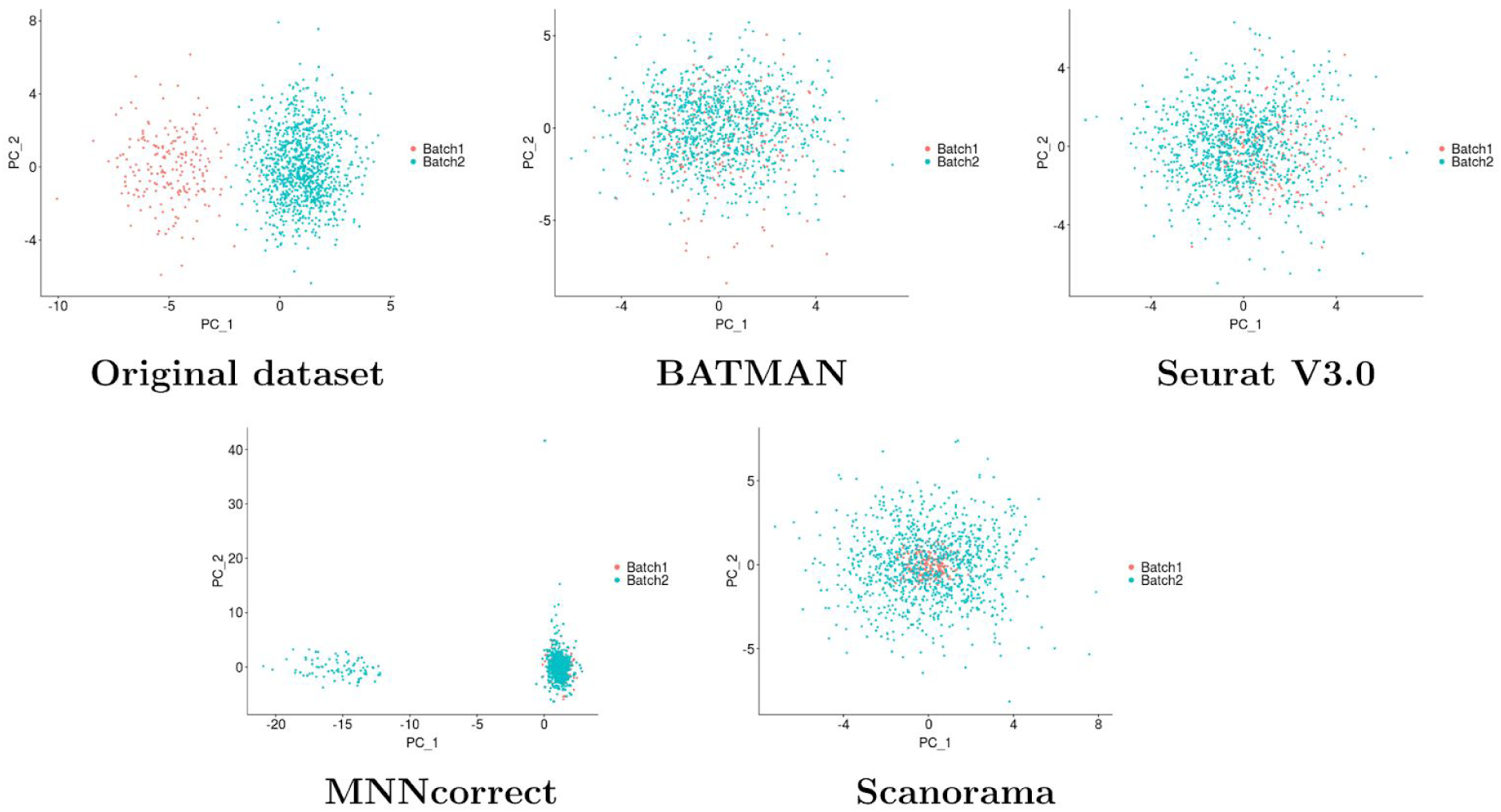
Simulated datasets with unequal batch sizes (LB-UB scenario) - PCA plots. Each dataset consists of 1000 cells and 1000 genes. The top 2 PCs are plotted.

**Table S2:**
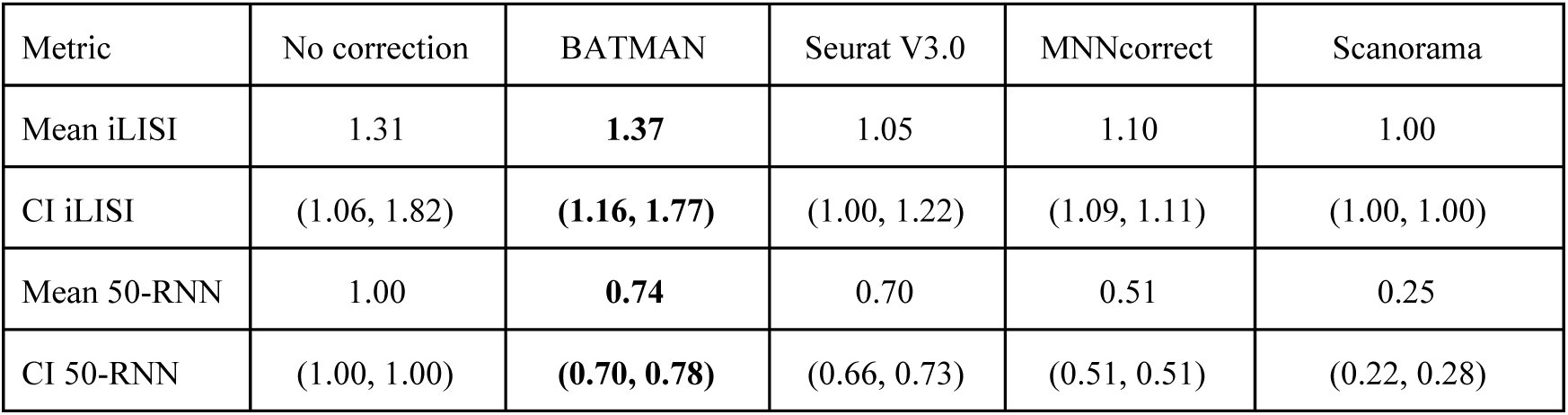
Integration results in LB-UB scenario (large batch effects, unequal batch sizes): iLISI and 50-RNNscores. The best results are emphasized in bold. CI means confidence interval.

| Metric | No correction | BATMAN | Seurat V3.0 | MNNcorrect | Scanorama |
| --- | --- | --- | --- | --- | --- |
| Mean iLISI | 1.31 | **1.37** | 1.05 | 1.10 | 1.00 |
| CI iLISI | (1.06, 1.82) | **(1.16, 1.77)** | (1.00, 1.22) | (1.09, 1.11) | (1.00, 1.00) |
| Mean 50-RNN | 1.00 | **0.74** | 0.70 | 0.51 | 0.25 |
| CI 50-RNN | (1.00, 1.00) | **(0.70, 0.78)** | (0.66, 0.73) | (0.51, 0.51) | (0.22, 0.28) |

**Figure S4:**
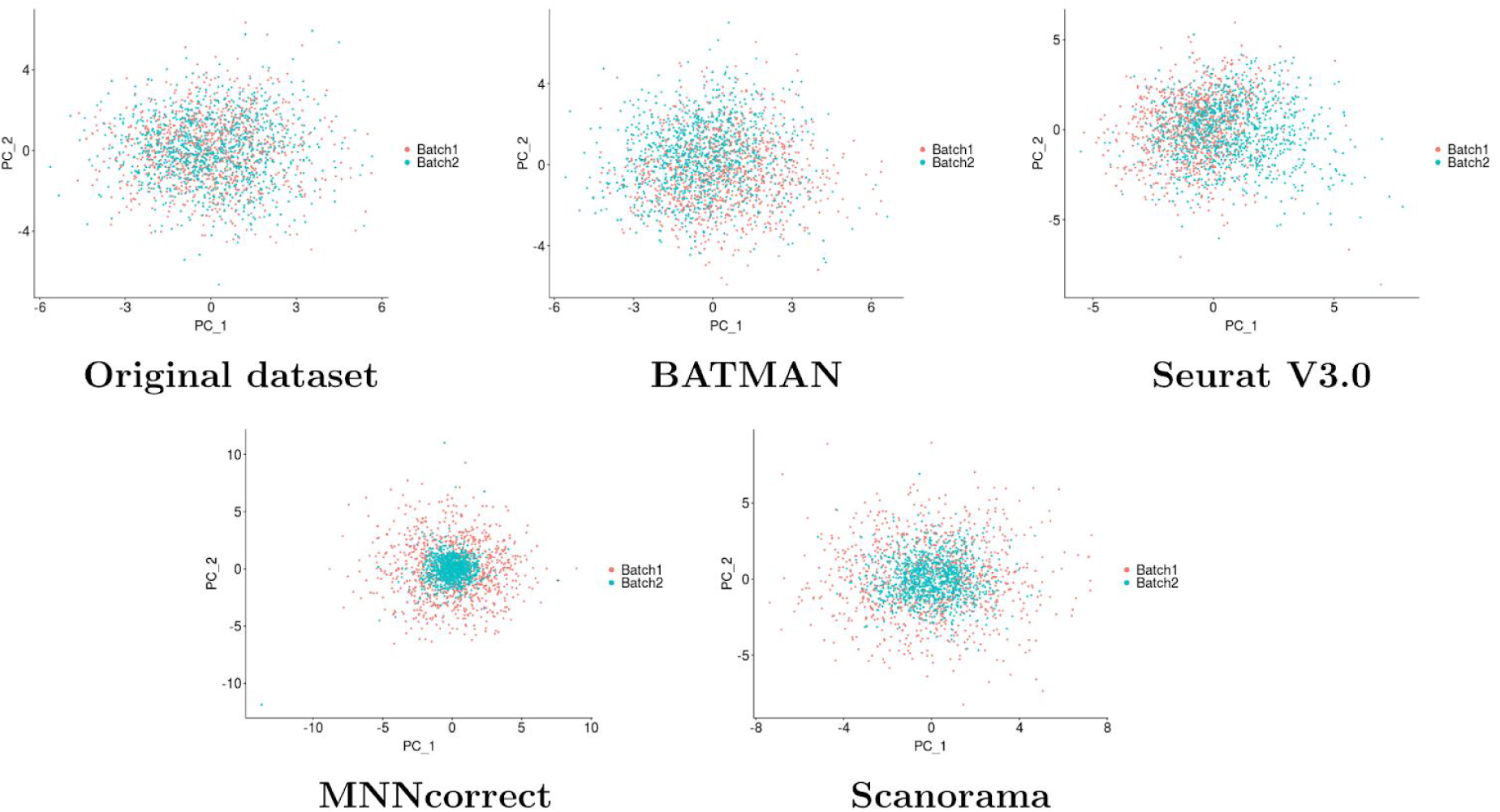
Simulated datasets with small batch sizes (SB scenario) - PCA plots. Each dataset consists of 1000 cells and 1000 genes. The top 2 PCs are plotted.

**Table S3:** Integration results in SB scenario (small batch effects, unequal batch sizes): iLISI and 50-RNNscores. The best results are emphasized in bold. CI means confidence interval.

| Metric | Original | BATMAN | Seurat V3.0 | MNNcorrect | Scanorama |
| --- | --- | --- | --- | --- | --- |
| Mean iLISI | 1.88 | **1.87** | 1.00 | 1.00 | 1.00 |
| CI iLISI | (1.70, 1.94) | **(1.66, 1.93)** | (1.00, 1.01) | (1.00, 1.00) | (1.00, 1.01) |
| Mean 50-RNN | 1.00 | **0.93** | 0.58 | 0.51 | 0.13 |
| CI 50-RNN | (1.00, 1.00) | **(0.91, 0.95)** | (0.56, 0.60) | (0.51, 0.52) | (0.11, 0.16) |

**Figure S5:**
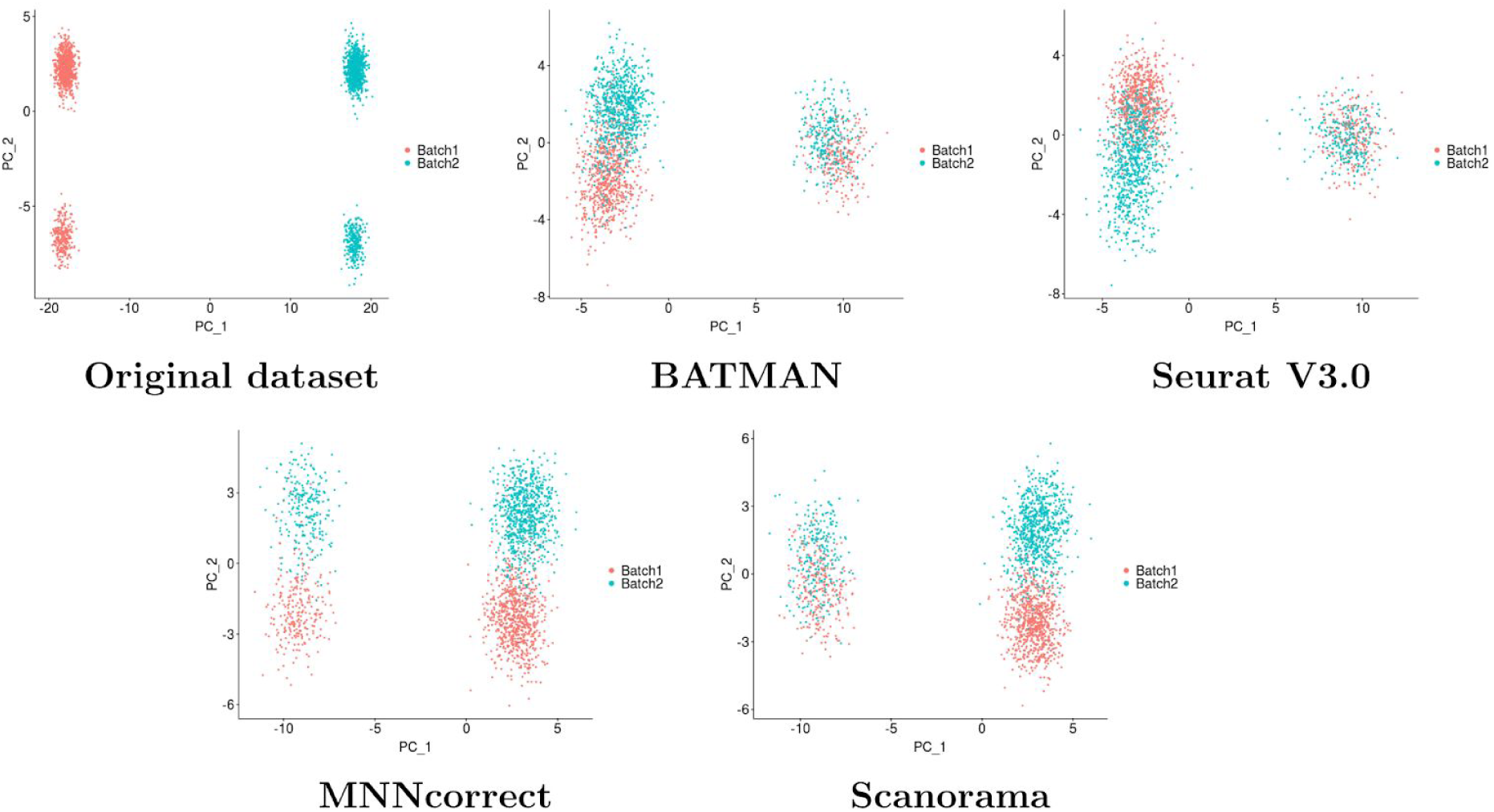
Simulated datasets with large batch effects and two cell types (LB-CT scenario) - PCA plots. Each dataset consists of 1000 cells and 1000 genes. Cell type frequencies are 80% and 20%. The top 2 PCs are plotted.

**Table S4:** Integration results in LB-CT scenario (large batch effects, two cell types): iLISI and 50-RNN scores. The best results are emphasized in bold. CI means confidence interval.

| Metric | Original | BATMAN | Seurat V3.0 | MNNcorrect | Scanorama |
| --- | --- | --- | --- | --- | --- |
| Mean iLISI | 1.00 | 1.79 | 1.00 | **1.85** | 1.61 |
| CI iLISI | (1.00, 1.00) | (1.60, 1.87) | (1.00, 1.00) | **(1.85, 1.85)** | (1.61, 1.61) |
| Mean cLISI | 1.01 | 1.04 | 1.03 | **1.00** | **1.00** |
| CI cLISI | (1.01, 1.01) | (1.00, 1.22) | (1.03, 1.03) | **(1.00, 1.00)** | (1.00, 1.00) |
| Mean 50-RNN | 1.00 | 0.95 | 0.64 | **0.99** | 0.96 |
| CI 50-RNN | (1.00, 1.00) | (0.89, 0.98) | (0.64, 0.64) | (0.99, 0.99) | (0.96, 0.96) |

**Figure S6:**
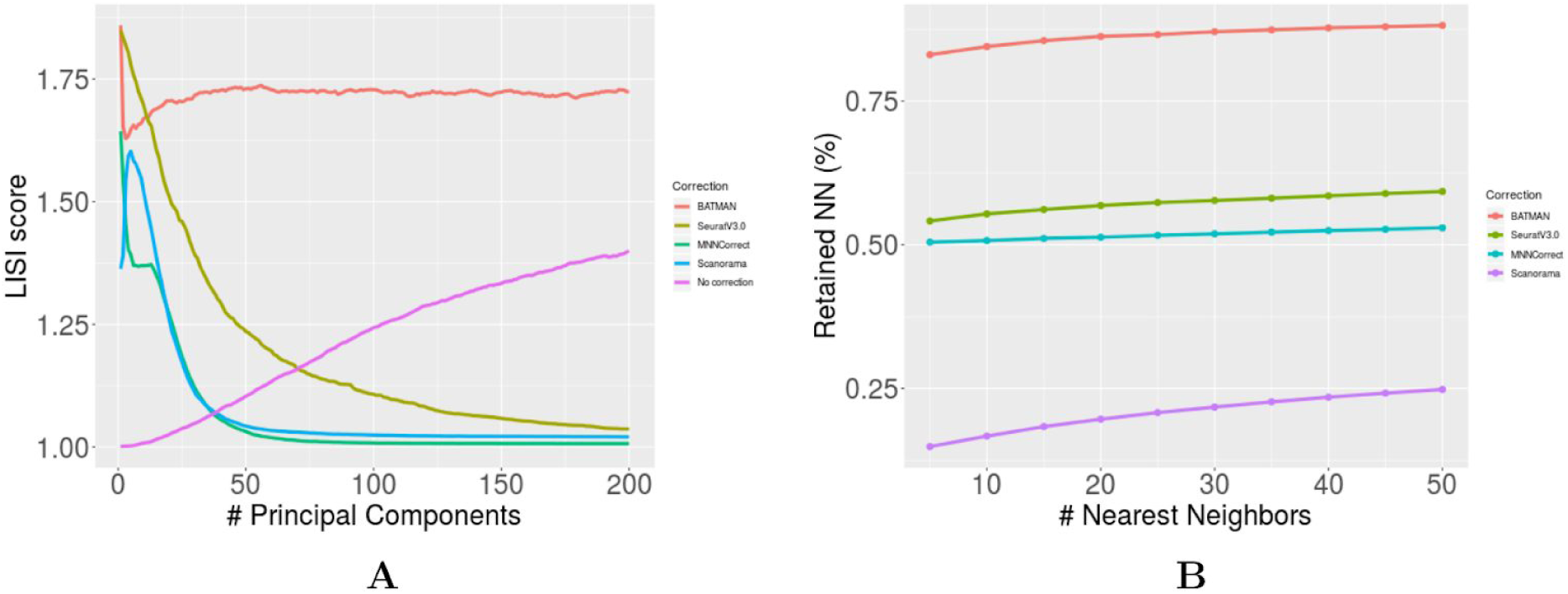
Evaluation of integration with large batch effects and large dropout (LB-DR scenario). A) iLISI score as a function of the number of top principal components. B) *k* -RNN score for different values of *k*.

**Figure S7:**
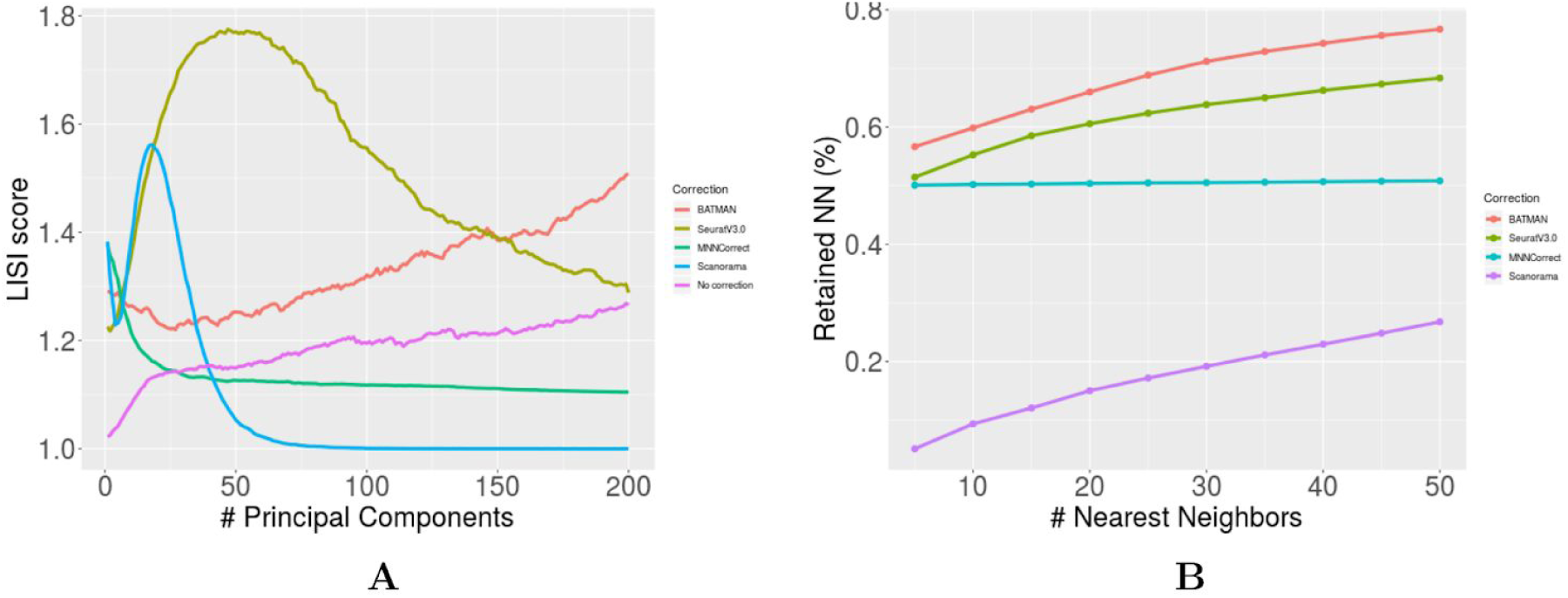
Evaluation of integration with large batch effects and unequal batch sizes (LB-UB scenario). A) iLISI score as a function of the number of top principal components. B) *k* -RNN score for different values of *k*.

**Figure S8:**
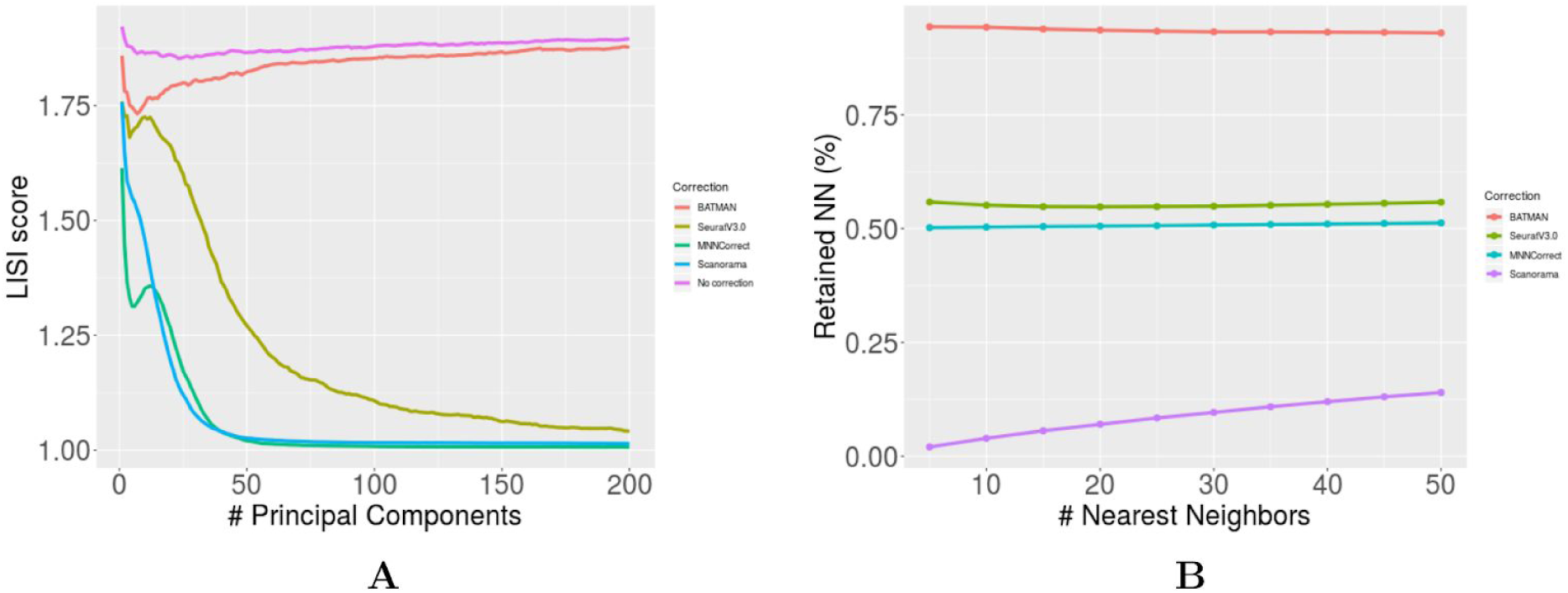
Evaluation of integration with small batch effects (SB scenario). A) iLISI score as a function of the number of top principal components. B) *k* -RNN score for different values of *k*.

**Figure S9:**
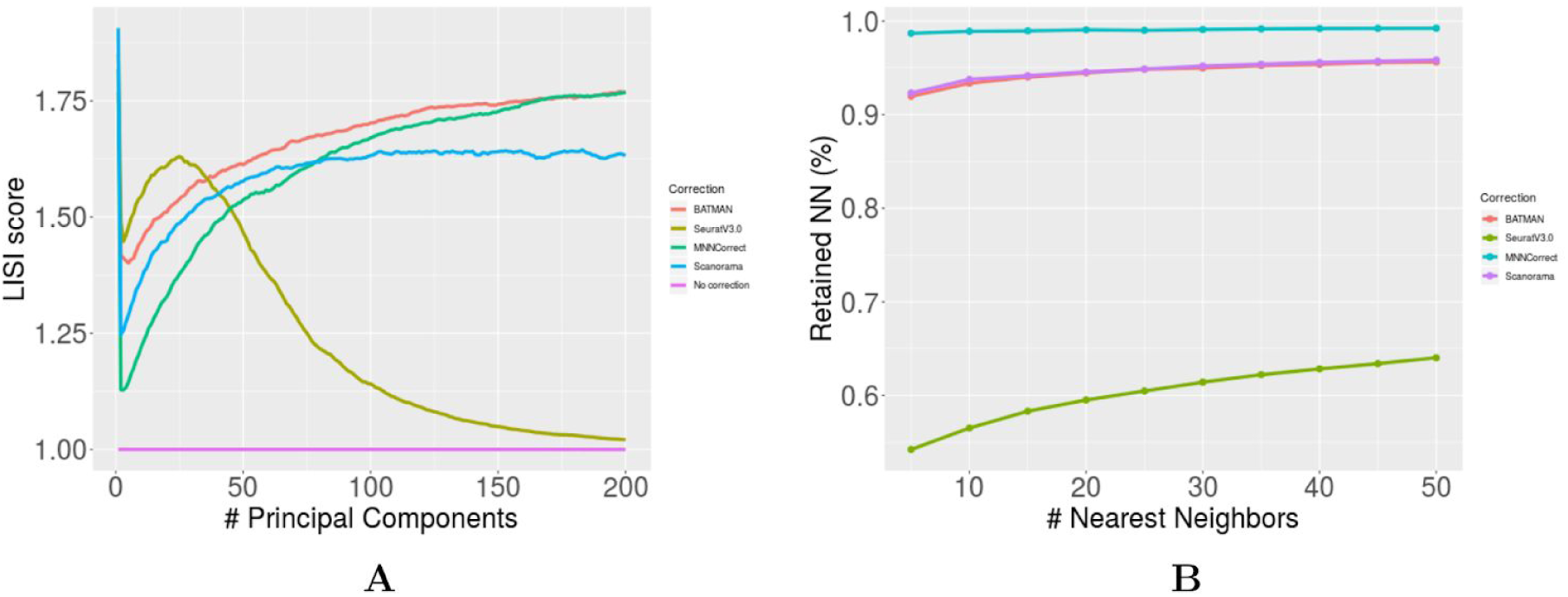
Evaluation of integration with large batch effects and two cell types (LB-CT scenario). A) iLISI score as a function of the number of top principal components. B) *k* -RNN score for different values of *k*.

**Figure S10:**
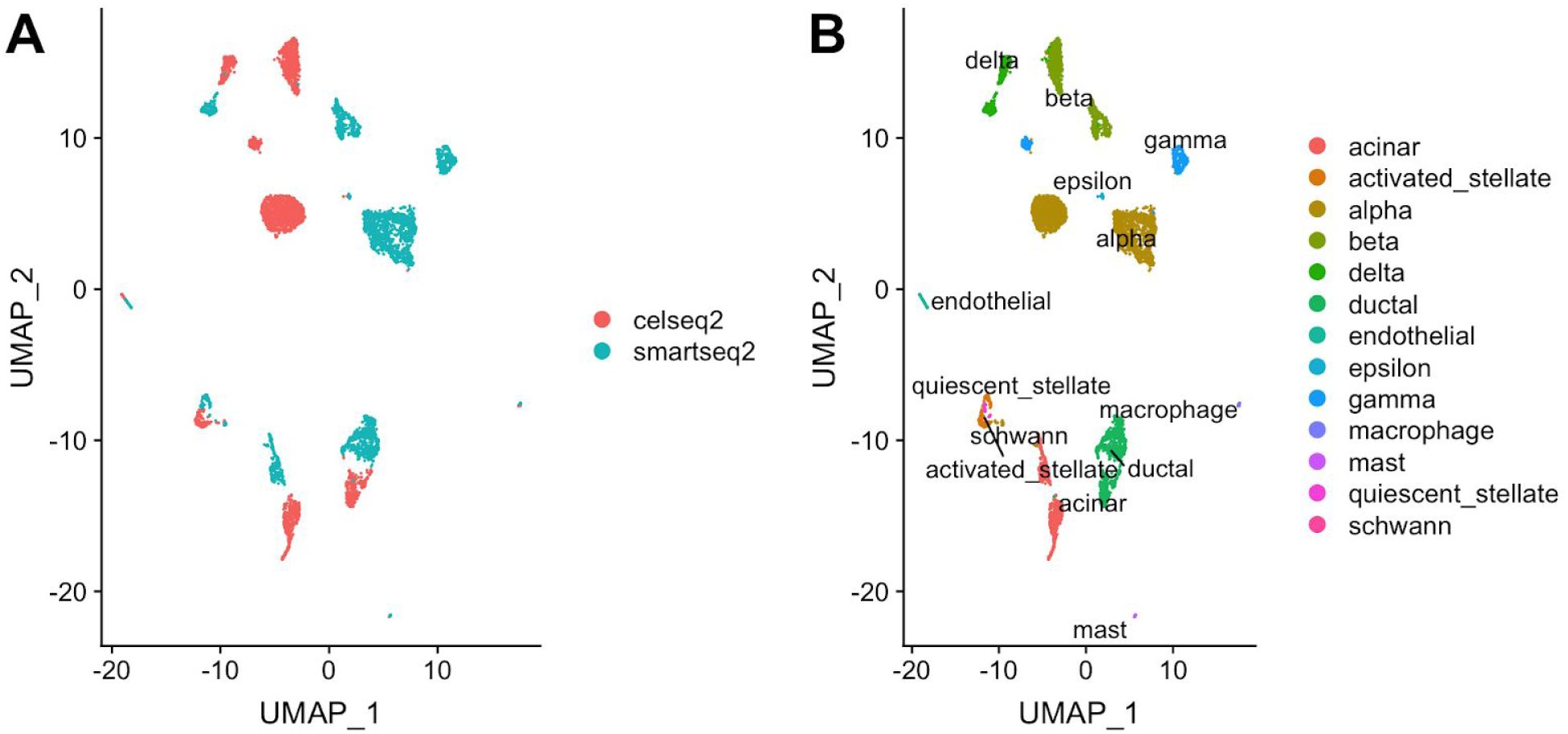
UMAP plots of two pancreatic datasets - CEL-Seq2 and Smart-Seq2. A) Datasets are colored by batch label; B) Datasets are colored by cell types.

**Table S5:** Cell-type composition of the two pancreas datasets (in %).

| Dataset | aci<br>nar | stell<br>ate | alpha | bet<br>a | delta | ductal | endo | epsil<br>on | gam<br>ma | mac<br>ro | mast | q-st<br>ella<br>te | sch<br>wan<br>n |
| --- | --- | --- | --- | --- | --- | --- | --- | --- | --- | --- | --- | --- | --- |
| CEL-Seq2 | 12 | 4 | 37 | 19.5 | 9 | 11 | 1 | 0.2 | 4.8 | 0.7 | 0.3 | 0.5 | 0.2 |
| Smart-Seq2 | 8 | 2.3 | 42 | 13 | 5.3 | 18.5 | 0.9 | 0.3 | 9 | 0.3 | 0.3 | 0.3 | 0.1 |

**Figure S11:**
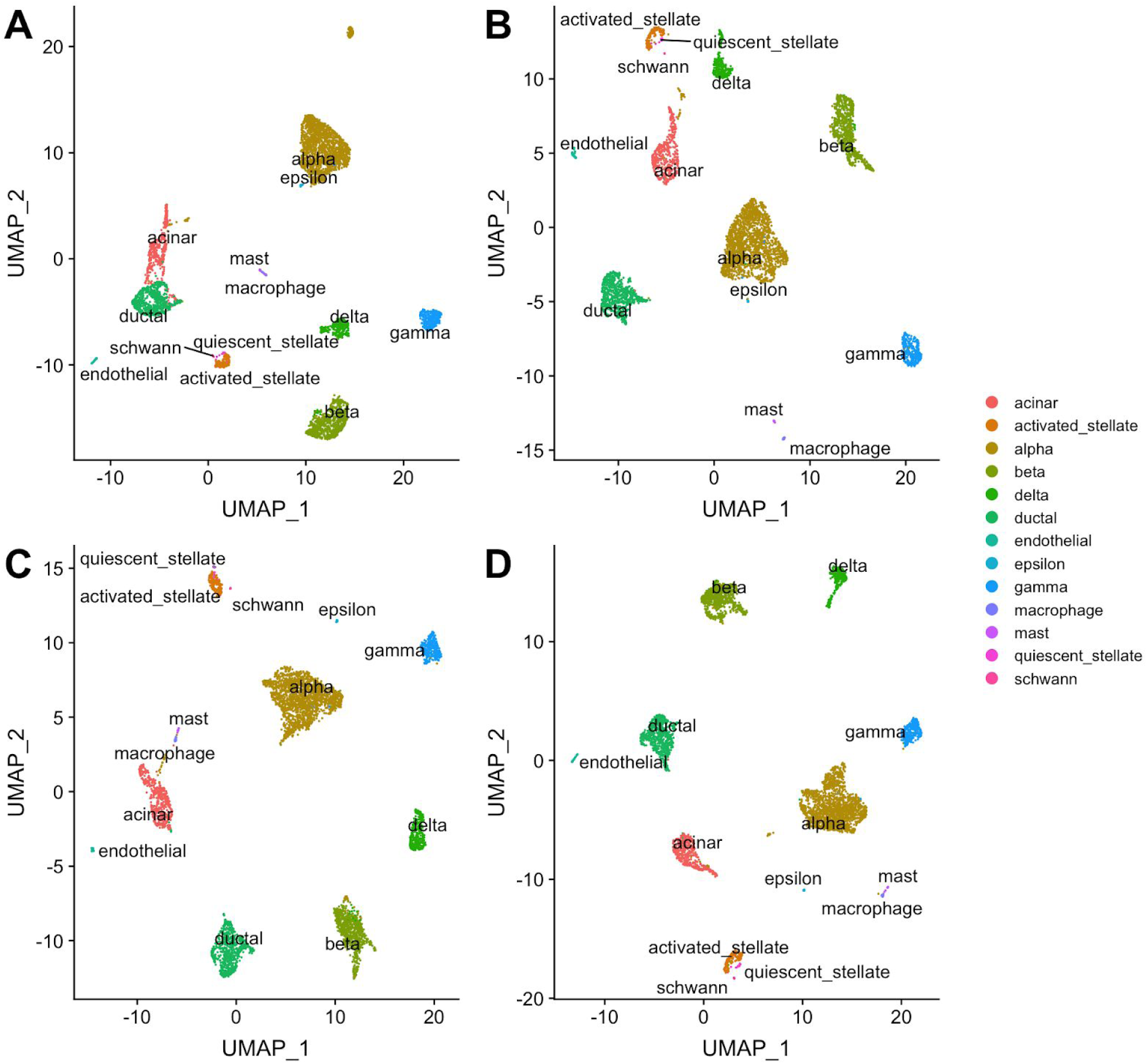
UMAP plots of two pancreatic datasets - CelSeq2 and SmartSeq2: integration results in “all cell types” experiment. Datasets are colored by cell types. A) BATMAN; B) Seurat V3.0; C) MNNcorrect; D) Scanorama.

**Figure S12:**
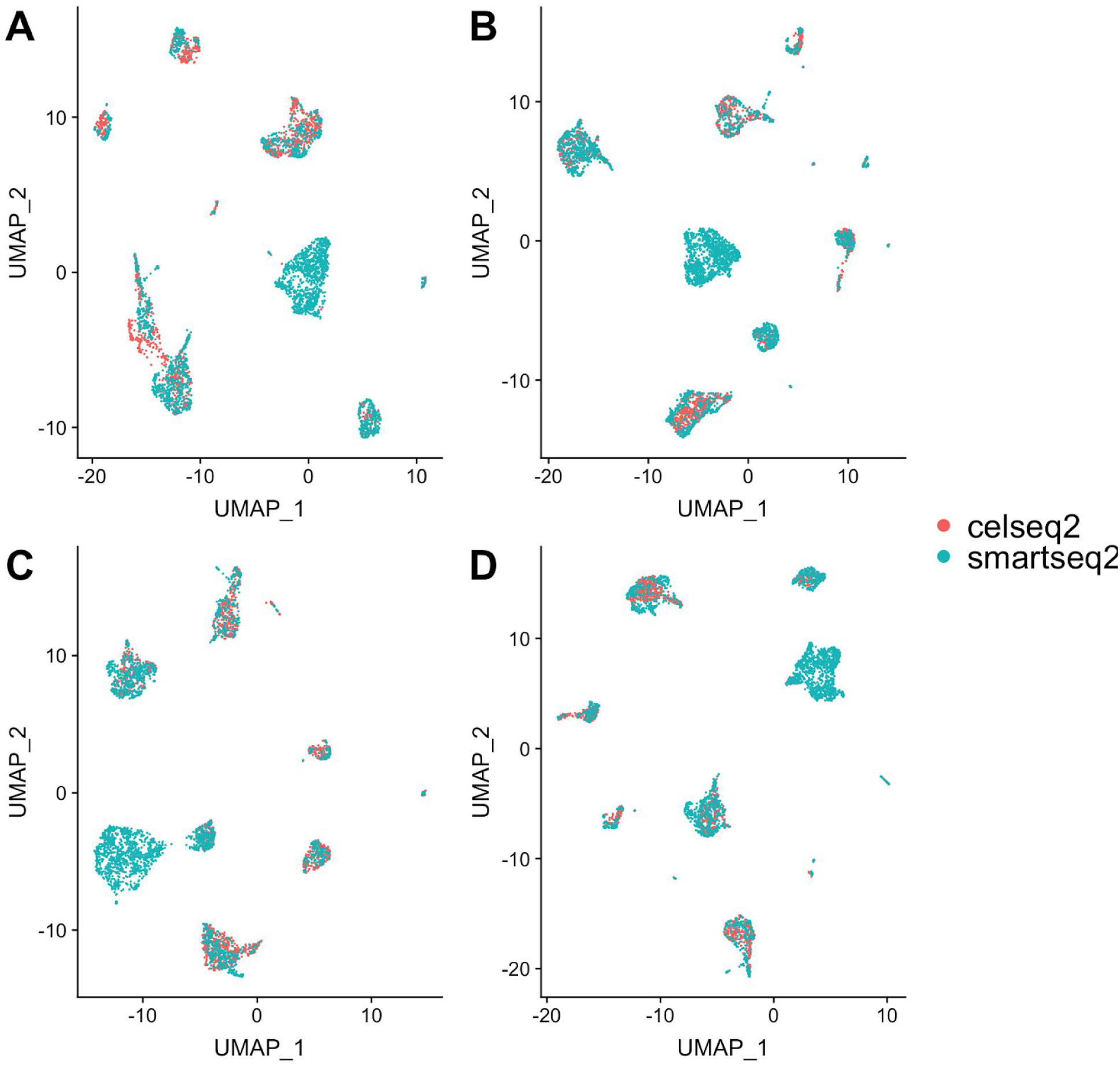
UMAP plots of two pancreatic datasets - CelSeq2 and SmartSeq2: integration results in “1-held out” experiment. Datasets are colored by batches. A) BATMAN; B) Seurat V3.0; C) MNNcorrect; D) Scanorama.

**Figure S13:**
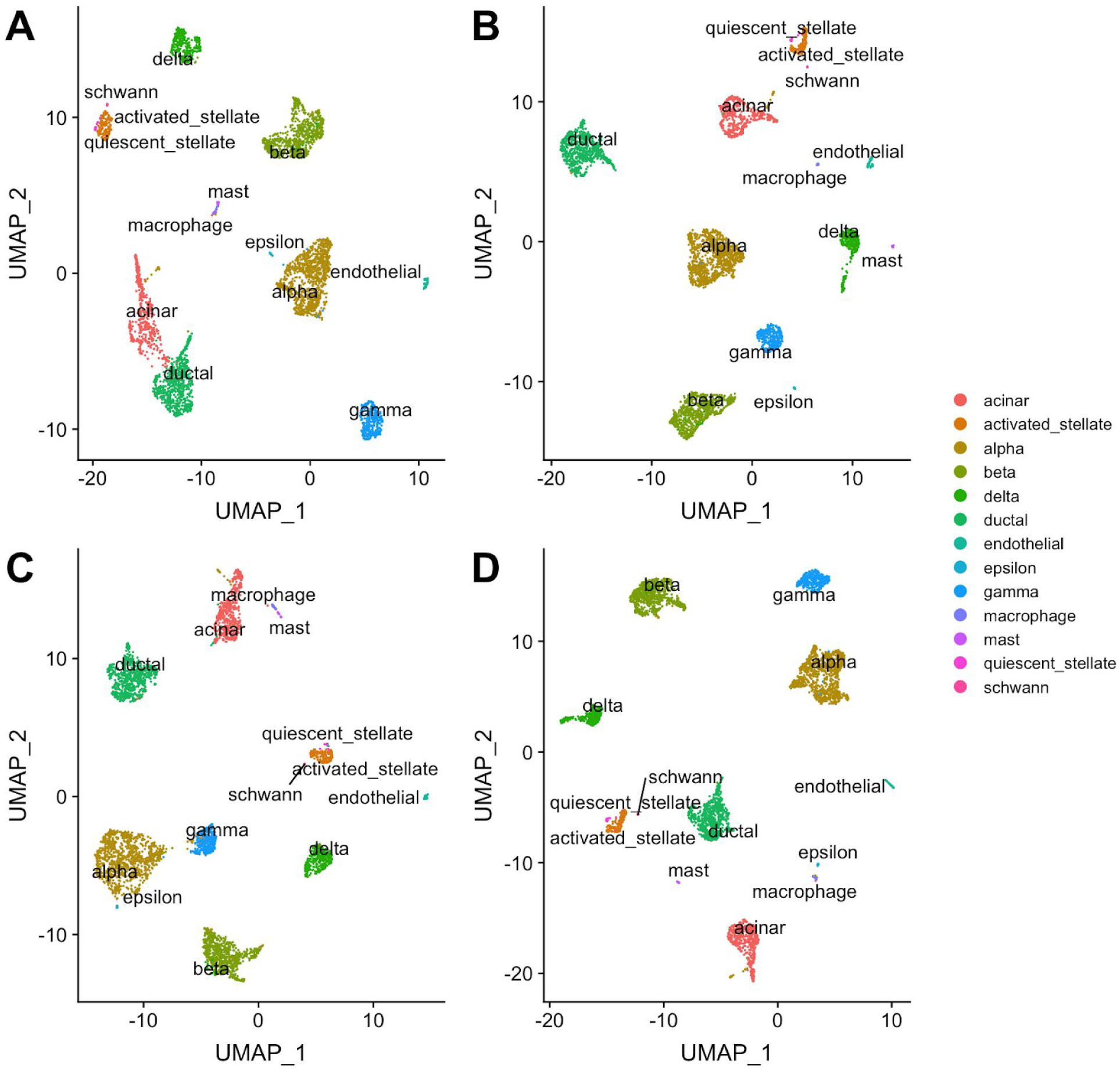
UMAP plots of two pancreatic datasets - CelSeq2 and SmartSeq2: integration results in “1-held-out” experiment. Datasets are colored by cell types. A) BATMAN; B) Seurat V3.0; C) MNNcorrect; D) Scanorama.

**Figure S14:**
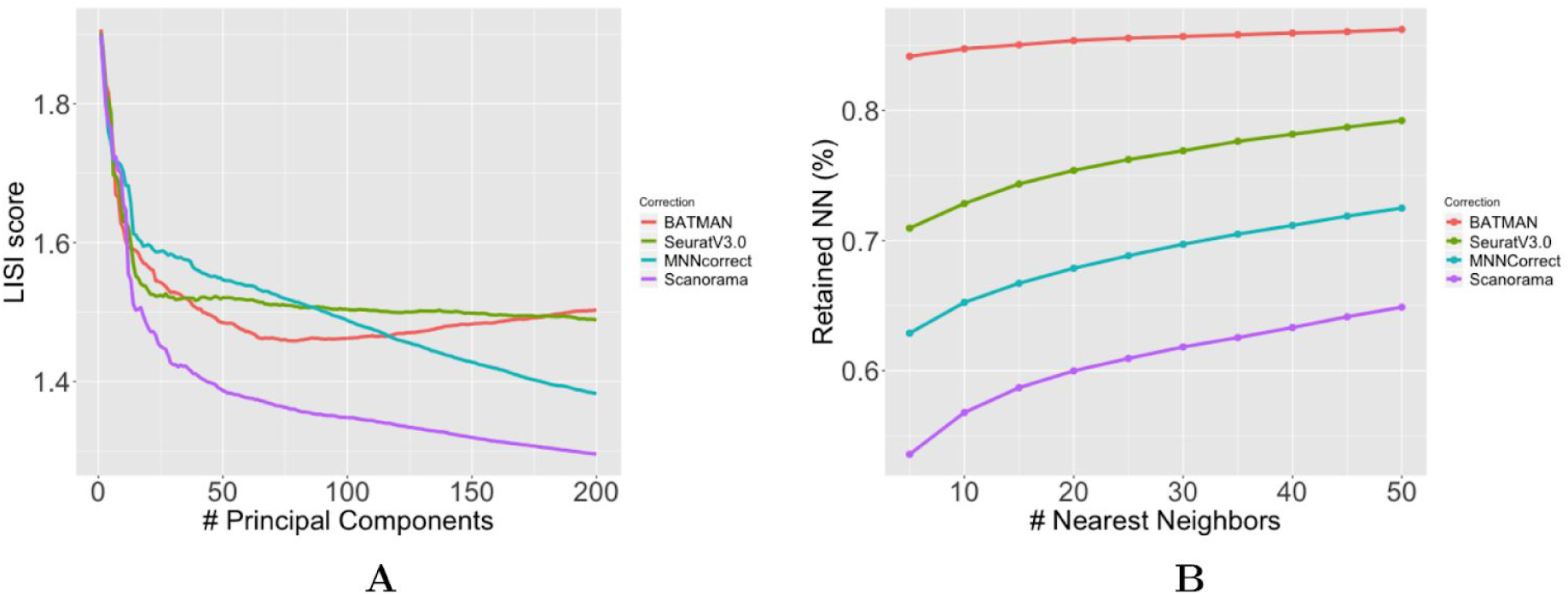
UMAP plots of two pancreatic datasets - CelSeq2 and SmartSeq2: integration results. Datasets are colored by cell types. A) BATMAN; B) Seurat V3.0; C) MNNcorrect; D) Scanorama.

